# RNA velocity in growing cells

**DOI:** 10.64898/2025.12.18.695252

**Authors:** Vishal Shah, Hia Ming, Brian Cleary

**Author notes:** Equal contribution (Vishal Shah, Hia Ming).

## Abstract

The ultimate promise of single cell “RNA velocity” methods is compelling: in principle, one can project forward the transcriptional state of each cell and map long-term expression trajectories. While there has been robust and ongoing articulation of limitations of existing methods, consensus frameworks fail to account for a fundamental aspect of cellular dynamics: growth. In a growing population, biomass (including RNA and other macromolecules) is constantly accumulating. This implies a homeostatic velocity (defined in the terms of production and degradation) that is positive, which is at odds with the conventional estimation, interpretation, and uses of velocity. Here, we investigate the consequences of omitting cell growth from the RNA velocity framework. We demonstrate systematic errors that arise in simulations of growing cells and show evidence for these artifacts in existing data. Finally, we point the way forward and highlight that explicitly accounting for cell growth can lead to new biological insights.

## Introduction

RNA abundance can be tightly regulated through control of kinetic rates of production, processing, and degradation. While transcriptional kinetics have been studied for many years, it has only recently become possible to study these processes with single cell resolution at scale in routine experiments^1–6^. New methods estimate single cell “RNA velocity” vectors, which capture the rate of change of each measured gene in a cell. Studying velocity at the single cell level allows one to access variability through natural heterogeneity (*e.g.*, observing cells in different phases of the cell cycle or at different points along a developmental trajectory) or, in principle, through pooled screens with each cell receiving a different perturbation – in either case enabling insight into the regulation of expression. Going further, a number of studies use velocity vectors to “project forward” the transcriptional state of each cell^7–20^. The ultimate promise of this approach is that, in observing velocity vectors of many single cells in a population, one can, in principle, piece together observed short-term changes in each cell and map long-term expression trajectories that were never directly observed, charting the paths that single cells take in dynamic processes and providing a foundation for understanding the phenotypic space that cells can occupy.

At a high level, there are two approaches to estimating RNA velocity. Early single cell RNA velocity profiling methods examined the ratio of spliced (older transcript) to unspliced (newer transcript) counts for each mRNA in each cell to infer production, splicing, and degradation rates, and to calculate single-cell velocity profiles^6,7^. More recent methods incorporate “metabolic” labeling with a nucleotide analog (*e.g.*, 4-thiouridine, 4sU) to infer rates by comparing unlabeled to labeled transcripts produced during a defined window^21–23^. A host of computational RNA velocity tools have been developed in parallel, applying various approaches to estimate the basic kinetic parameters^24^, and for downstream tasks like trajectory inference^9^ or RNA vector field mapping^8^.

While the ultimate promise of these methods is compelling, their limitations have also been widely documented^25–27^. Splicing based methods in particular suffer from substantial technical noise and bias (arising in part from the necessary but incidental capture of introns)^8^, as well as fundamental modeling and computational challenges^27^. Labeling based methods are generally believed to be significantly more robust, both because they do not rely on incidental capture and because labeling happens in a defined temporal window, allowing for observation of counts of transcripts produced or degraded in defined time periods.

Despite the robust and ongoing articulation of limitations and mitigation strategies, consensus RNA velocity frameworks have yet to account for a fundamental aspect of cellular dynamics: cell growth. Cell growth in dividing cells is known to be a global driver of RNA dynamics^28–32^, and though the majority of RNA velocity studies have been conducted in dividing cells, none seem to account for growth. We postulated that the omission of cell growth from RNA velocity frameworks leads to significant errors in estimation and interpretation. Here, we investigate those effects, focusing on labeling based strategies for velocity estimation, as these have emerged as the more quantitatively viable alternative.

## Results

### The basic setup: estimation and interpretation of RNA velocity

RNA velocity captures the rate of change for each measured gene. In the basic setup, the velocity for gene *i* is given by the balance of RNA production and degradation: *v_i_* = *α_i_* − *γ_i_x_i_*, where *α* is the production rate, *γ* the degradation rate, and *x* the gene abundance. (Here, “gene abundance” refers to the number of RNA molecules of each gene in the cell; we address alternative units – relative abundance and concentration – below). Calculation of velocity in this setup therefore requires that we observe gene abundance and estimate the two kinetic parameters. In single cell experiments, a common approach is to first estimate the population average degradation rate for each gene, then use those values together with observations of transcript production in a defined window to estimate production rates at the single cell level, and finally combine the terms to give single cell estimates of the velocity of each gene^3,8^ (**Figure 1A-B**).

**Figure 1:**
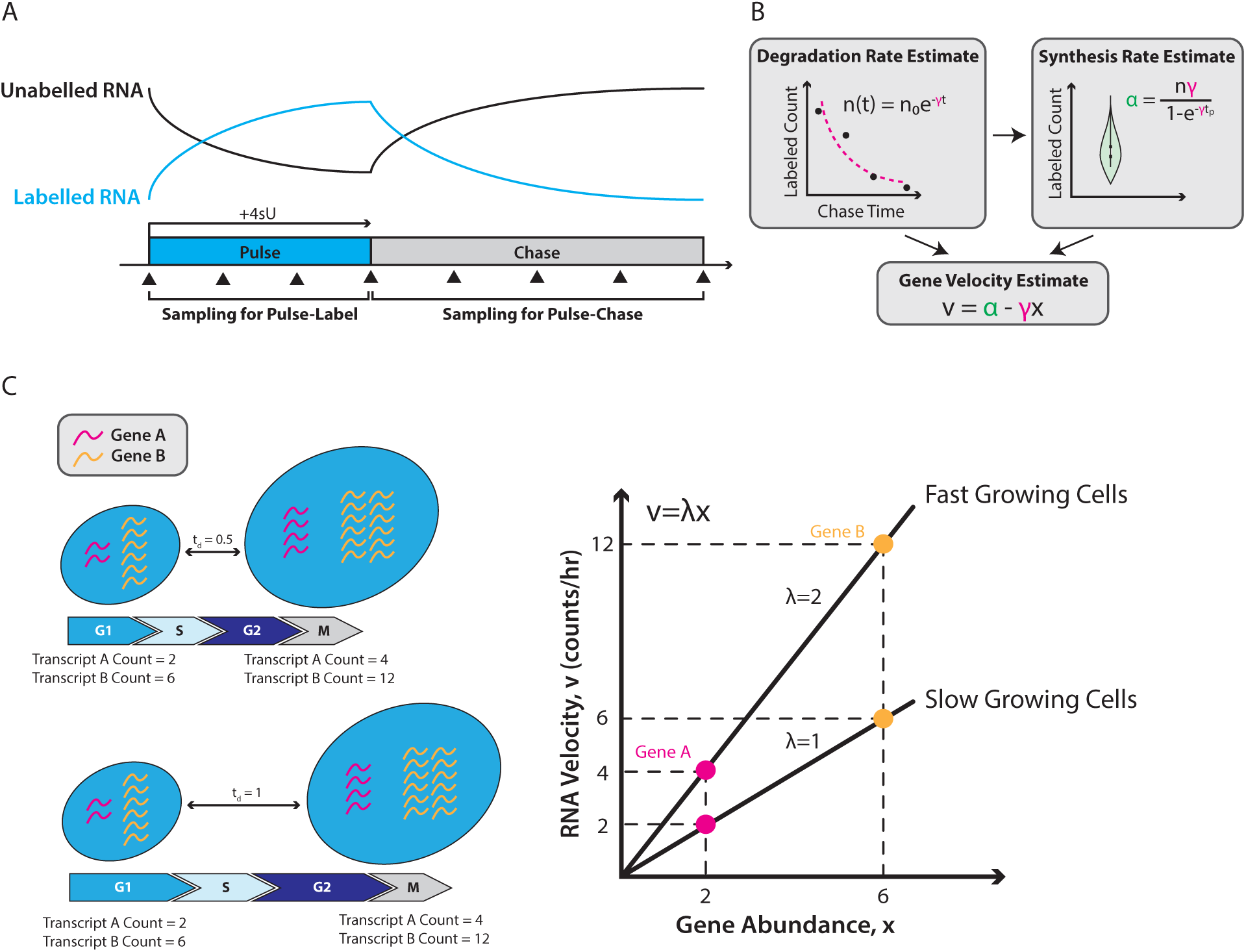
RNA velocity is proportionate to the growth rate of the cell. (A) Metabolic labeling-based RNA velocity experiments are typically of the pulse-label type or the pulse-chase type. In the pulse-label experiment, samples are taken while cells are being exposed to the metabolic label (e.g. 4sU) - i.e. the ‘pulse’; in the pulse-chase experiment, samples are taken after a defined pulse duration in the absence of the metabolic label, usually while exposed to high levels of uridine - i.e. the ‘chase’. The typical profile of RNA labeled or unlabeled with the metabolic label is shown for the duration of the pulse and chase. (B) Gene-specific RNA degradation rate (*γ*) is estimated by fitting an exponential decay function to the mean abundance of labeled RNA (n) sampled from the chase. Gene-specific RNA synthesis rate (*α*) is estimated at a single cell level from labeled RNA abundances measured after a defined pulse duration by incorporating the previously calculated cell-averaged degradation rate. A gene- and cell-specific RNA velocity (v) is estimated from the calculated degradation and synthesis rates, as well as the total (labeled and unlabeled) RNA abundance (x) of the gene. (C) Relationship between RNA velocity and gene abundance for fast and slow growing cells. Fast growing cells with a growth rate λ=2 (doubling time t_d=0.5) have a higher homeostatic RNA velocity for the same steady-state gene abundance as slower growing cells.

These velocity estimates are interpreted in relation to a steady state: velocity of zero is presumed to be at steady state, with positive and negative velocities reflecting ongoing up- or down-regulation. In downstream applications, velocities are used to project forward to the future transcriptional state of each cell. In the ideal scenario, these short-term projections across a population of cells can be stitched together, allowing for the reconstruction of long-term trajectories that were never directly observed^8,9,11–20^.

While there are a number of models with elaborations on this framework, they all build on the same basic setting, which has unfortunately left out a key aspect of cellular dynamics: growth. In a growing population, biomass (including RNA and other macromolecules of the cell) is constantly accumulating. This is true too at the single cell level: biomass accumulates from the beginning of cell cycle to the end before division brings daughter cells roughly back to the same size and state (**Figure 1C**, left). This implies that to keep up with cell growth we expect a homeostatic velocity (defined in the usual terms of production and degradation) that is positive, which appears to be at odds with the interpretation and uses of velocity outlined above.

This gap is especially important to examine in light of the widespread adoption of velocity methods. From a survey of 43 publications that develop, benchmark, or review RNA velocity methods and that collectively represent major methodological foundations of the field (**Supplementary Table 1**), we find that 40/43 did not address growth rates at all. The remaining 3/43 only partially dealt with some aspect of growth but made no correction for it when estimating RNA velocity. These 43 studies have been collectively cited in 4,836 unique publications with RNA velocity analysis; in a more examined subsample nearly all (∼98%) appear to do so in proliferative cells (**Methods**).

The following sections of this paper investigate the consequences of omitting cell growth from the RNA velocity framework. We first provide a brief review of the literature on RNA and protein dynamics in growing cells, then, through simulation, demonstrate systematic errors in interpretation and estimation that arise from ignoring cell growth. We show how inefficient detection and sampling similarly give rise to systematic artifacts in growing cells. In re-analysis of existing datasets, we observe the same biases present in simulation. Finally, we point the way forward for correcting some of these issues and highlight that explicitly accounting for cell growth in the RNA velocity framework can lead to new biological insights.

### Overview of Simulation Framework

In the course of the cell cycle, cells grow to approximately double in volume before dividing. A number of experiments have shown that both protein and RNA content scales with cell volume, in cycling and non-cycling cells, to maintain approximately constant concentration. For proteins, prior work has shown that protein numbers, the rate of change of protein mass, and the rate of protein production are proportional to cell volume and mass^33–37^. Similar results have been found for mRNA in yeast and mammalian cells^30,38–40^, with work in mammalian cells showing that the copy number of individual genes and the production rate correlate with cell volume^28,38–41^ and that total mRNA counts increase and double across the cell cycle^28,41^.

At a high level, these results can be captured with a simple model: mRNA abundance will double over the cell cycle, so that when the cell divides, with each daughter cell receiving approximately half the RNA content of the parent, the mRNA abundance of each daughter cell will return to the original abundance, maintaining a steady state. This gives rise to a positive correlation between the abundance of a gene and its velocity (**Figure 1C**). A gene with a starting abundance of 1 copy will need to produce 1 additional copy by the end of the cycle. A second gene with a starting abundance of 10 will need to produce 10 additional copies in the same time frame. Thus, velocities are proportional to abundance, with a slope given by growth rate. Up- or down-regulation would manifest as deviations from this line.

A recently developed mathematical framework captures this idea in a more sophisticated, yet still relatively simple model with alternative scenarios given by different limiting factors^42,43^. In the setting most consistent with available data in exponentially growing cells, cell growth and protein production are ribosome limited, and mRNA production is limited by RNA polymerase. In this case, RNA and protein production rates both scale with cell volume, such that transcription rates increase exponentially across the cell cycle. Degradation rates are assumed to be constant for simplicity, though some evidence suggests these too might adjust (decrease) across the cell cycle to maintain homeostasis^38,44,45^. Other scenarios in which RNAP are not limiting are possible as well. For example, if DNA templates were limiting, then the production rate of RNA transcripts may be constant across the cell cycle. Regardless, in each scenario RNA abundance must on average double before cell division.

We developed a simulation framework to systematically evaluate the effects of (unaccounted) cell growth on estimation and interpretation using this basic RNA velocity setup (**Figure 2**). Similar to the simulations developed alongside the above mathematical framework mentioned above^42^, we used the stochastic simulation algorithm (SSA) to simulate mRNA production, degradation, and cell growth and division in single cells, tracking a growing population of cells over time (**Figure 2A-C; Methods**). On top of these ground truth dynamics, we also included the (potentially noisy) experimental process of RNA labeling and observation (**Methods**).

**Figure 2:**
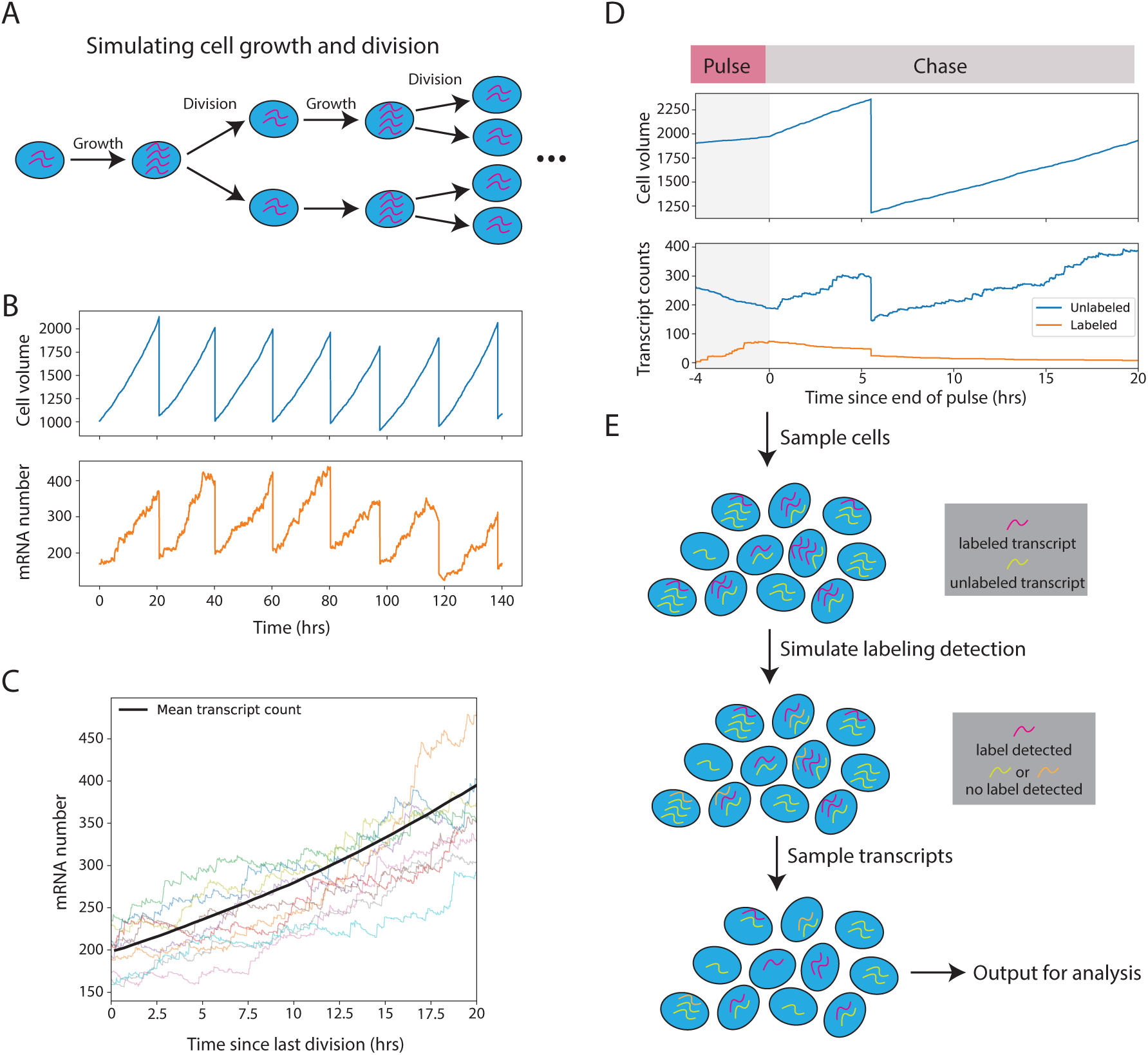
Simulation framework for modeling RNA velocity experiments in dividing cells (A) Schematic of the stochastic simulation framework. Cells double their volume and divide, partitioning transcripts equally between daughter cells. (B) Example simulation traces showing cell volume dynamics (top) and the abundance trajectory of a representative gene (bottom) across successive cell cycles. (C) Mean simulated transcript counts (black) for the gene in (B) across many cells (individual colored lines). (D) Diagram depicting cell volumes (top) and labeled/unlabeled abundances for the gene in (B) (bottom) in a pulse-chase experiment. This cell is slowed during the pulse, simulating 4sU toxicity. (E) Overview of the sampling and detection pipeline. For each simulated cell, labeled and unlabeled transcripts are subjected to imperfect detection and capture efficiency to obtain observed counts (see Methods).

We focus our analysis on two common experimental setups: one pulse-label setting containing a 4-hour pulse period, and one pulse-chase setting containing a 4-hour pulse and a 20-hour chase (**Figure 1A**). Both experiments were sampled every hour, during the pulse in the former, and during the chase in the latter. For now, we present our analysis in terms of RNA copy number and will discuss the use of normalization and RNA concentration in a later section.

### Growth rate effects on RNA velocity

We first evaluate growth rate effects in a noiseless scenario, with perfect detection efficiency (*δ* = 1) and complete sampling of all transcripts (that is, zero technical noise, but still allowing biological noise). We performed simulations with cell doubling times of 12 hours (*λ* ≈ 0.058) and of 20 hours (*λ* ≈ 0.035), and in non-dividing cells (infinite doubling time). Below, we walk through the steps of basic RNA velocity estimation, starting with degradation rates, then production rates, velocity, and trajectory estimation.

### Degradation rate

When growth is not explicitly represented, the estimated RNA degradation rates are systematically biased, having absorbed a growth rate effect. Across genes, growth rates, and experimental designs, the estimated degradation rates are almost exactly *γ*^∗^ = *γ* + *λ* (**Figures 3A,B** and **S1A-C**).

**Figure 3:**
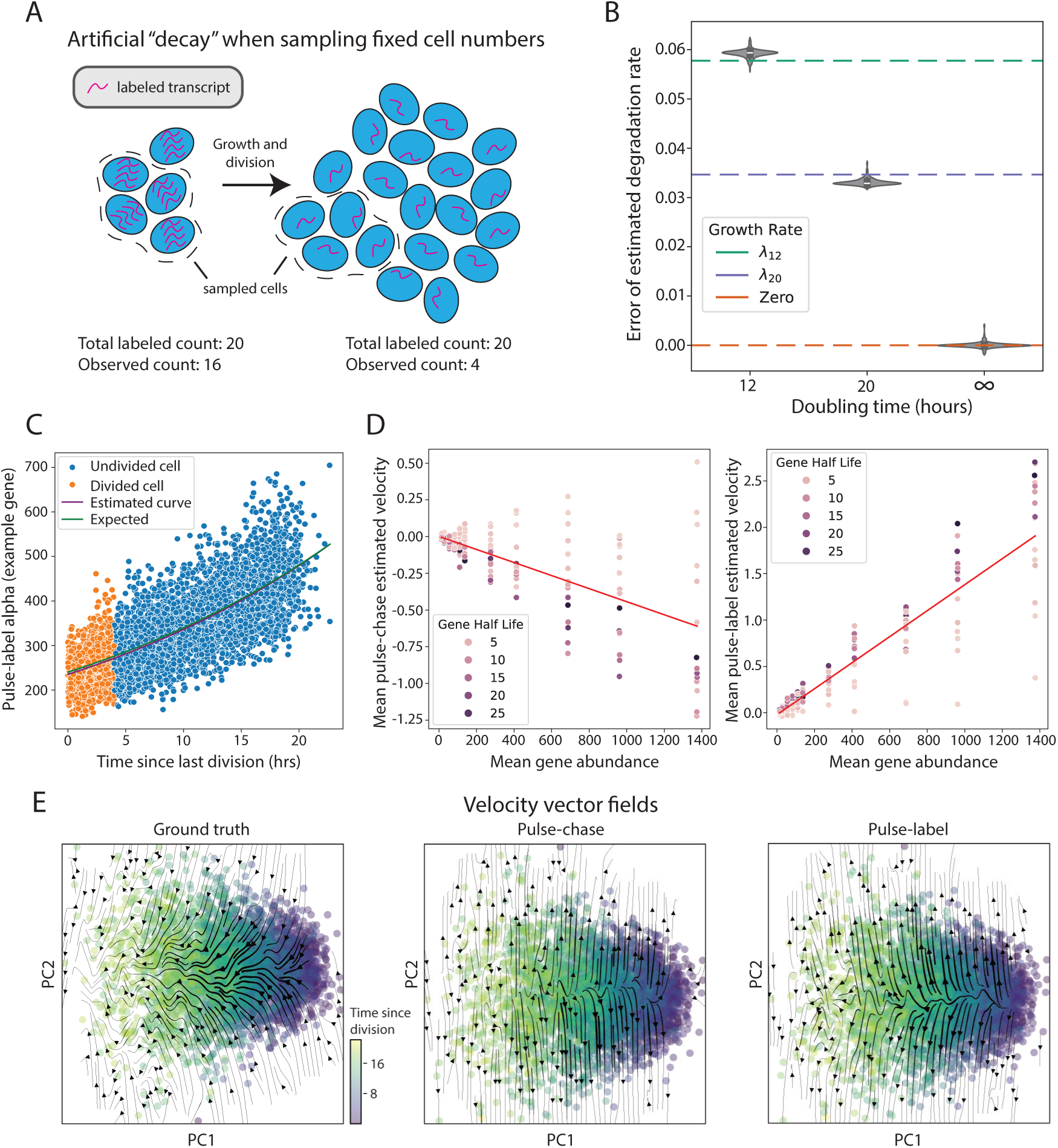
Growth rate is absorbed into degradation rate estimates with existing designs (A) Illustration of how sampling a fixed number of cells in a growing population leads to apparent (but artificial) transcript decay. In the example, the labeled transcripts for a given gene in a chase window have a degradation rate *γ*=0, and hence the total number of transcripts does not change from one timepoint to the next. (B) Violin plots showing the absolute error (y-axis) between true and estimated degradation rates for non-growing populations or with a 12- or 20-hour doubling time in a pulse-label experiment. Horizontal dashed lines show the error expected when any one of the three growth rates is absorbed into degradation estimates. (C) Scatter plot showing the estimated production rate (y-axis) for a single gene (*γ*=0.2057) in a population of cells (individual dots) as a function of each cell’s time since last division (x-axis). Colors for each dot indicate whether a cell underwent division between the start of the pulse and the time of sampling. Curves depict a best fit to the scatter plot (purple) and the expected production rate in exponentially growing cells (green). (D) Scatter plots of the population mean estimated velocity (y-axis) as a function of mean abundance (x-axis) for individual genes (dots) in pulse-chase (left) or pulse-label (right) experiments for cells with 20-hour doubling times. (E) Vector field plots generated from ground truth (left), pulse-chase (center), or pulse-label (right) velocity vectors for each cell (dots), projected into two dimensions (PC1 and PC2). Colorbar indicates each cell’s time since last division.

To illustrate why the growth rate effect is absorbed, consider a pulse-chase experiment and a gene with exceptionally long half-life (near zero degradation rate; **Figure 3A**). Labeled transcripts are produced during the pulse, then samples taken during the chase are meant to track decay as the RNA is degraded. With degradation near zero there is no actual decay, and the labeled transcripts persist through the course of the experiment. Nevertheless, there is a dilution of the transcripts at the single cell level (when a cell divides, each daughter receives approximately half the labeled transcripts). Since typical scRNA-Seq experiments capture a fixed number of cells per sample, rather than a fixed fraction – in this case, of a growing population – and the labeled content per cell is decreasing (even as the overall labeled content is unchanged), there is an apparent and artifactual “decay” of total labeled counts over time.

The contribution of cell growth and division to the inferred RNA decay rates have been recognized outside the RNA velocity literature in earlier studies^46,47^ and accounted for to some extent in in the broader literature of RNA kinetics^48–50^. However, current RNA velocity methods have yet to incorporate growth rate effects into their models (**Supplementary Table 1**). Under the current framework, growth rates are absorbed into the estimated degradation rate *γ*^∗^ and the primary effect of growth rate on degradation, therefore, is a misinterpretation of the parameter.

### Production rate

Although degradation rates are used in the estimation of the production rate *α*, production rates remain accurately estimated in the noiseless setting (**Figures 3C** and **S1D-E**). Recall, however, that production rates in the RNA polymerase-limited setting increase exponentially over the cell cycle, and that, unlike degradation rates, *α* is estimated at the single cell level. Thus, for a given gene in the absence of “interesting” (*i.e.*, non-homeostatic) regulatory variation, there will still be a two-fold variation in observed production rates. This again complicates interpretation; we return to this below after discussing additional effects.

### Velocity and trajectories

With accurate estimation of *α* and growth rate absorbed into *γ*^∗^, velocities do not have the velocity-abundance relationship expected if velocity is given by the balance of production and degradation (**Figure 1C**), and are closer to zero (**Figures 3D** and **S1F**). There remains, however, a small systematic bias in the abundance vs velocity plots that varies based on growth rate and experimental design.

Although the velocity bias due to growth effects appears small in the noiseless setting, it is nevertheless consequential for downstream analysis of single cell trajectories. We can see this by following the basic procedure for trajectory analysis with the velocity values from our simulated populations (**Methods**). In brief, we project the velocity vector for each cell into low dimensions, then use common tools to visualize smoothed trajectories.

First, we examine visualizations of ground truth velocity vectors. Since there are no ongoing regulatory changes other than homeostatic dynamics, we expect the only relevant trajectories to track cell cycle progression. Indeed, trajectory plots of ground truth velocity vectors show cells moving from right to left across PC1, in the direction of cell cycle progression (**Figure 3E**, left). In contrast, performing the same analysis with estimated velocity vectors produces coherent, diverging trajectories that are orthogonal to the true (and only salient) direction of transcriptional change (**Figure 3E**, middle and right).

Thus, while the absorption of growth rates into *γ*^∗^ pushes velocity estimates to nearly zero, potentially allowing for “correct” interpretation of substantially positive or negative velocities as reflecting non-homeostatic regulation, downstream trajectory analysis is sufficiently sensitive to remaining artifacts to potentially produce wildly misleading conclusions.

### Variable growth rate effects

Next, we consider what happens when growth rates vary across the duration of an experiment or across the population of sampled cells. For the former, we consider slowed growth during the pulse, which results from defects caused by introduction of the commonly used nucleotide analog 4sU. We then consider the case when two populations with different growth rates are mixed, and RNA velocity analysis is performed without accounting for variable growth.

### Growth defects associated with 4sU

Introduction of the nucleotide analog 4sU during the pulse window can slow, or even halt cell growth, as seen in previous reports^21,51^ and in our own data (**Figure S2A**). Effects from 4sU defects may manifest differently in pulse-label and pulse-chase experiments. In a pulse-label experiment samples are taken only during the pulse, and so all samples would be at the same – albeit non-physiological – growth rate. In a pulse-chase experiment, cells will grow at one (lower) rate during the pulse, then another during the chase. To capture these effects in simulation, we applied an 80-hour doubling time during the pulse window and a 20-hour doubling during the chase. As expected, estimated *γ*^∗^ from pulse-label experiments incorporate a growth rate associated with 80-hour doubling, while pulse-chase *γ*^∗^ values incorporate 20-hour doubling (**Figures 4A** and **S2B**). Although there is no difference in RNA degradation rates in these settings, the two experimental designs result in substantially different estimations of *γ*^∗^.

**Figure 4:**
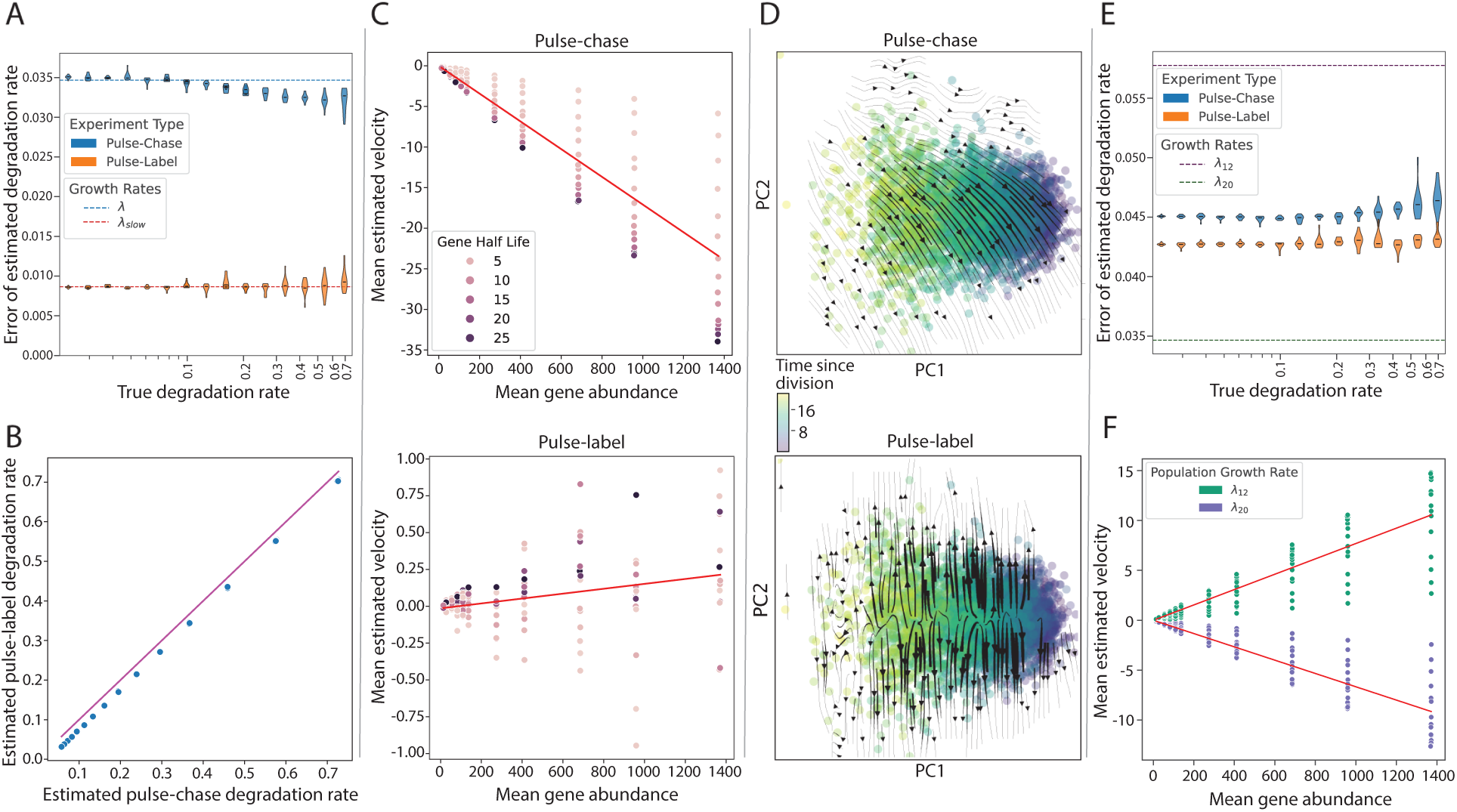
Variable growth rates result in inconsistent growth rate incorporation (A) Violin plots showing the absolute error (y-axis) between true and estimated degradation rates across genes under 4sU-induced slowdown. Errors are stratified by ground-truth degradation rates (x-axis) and shown for both pulse-chase (blue) and pulse-label (orange) experiments. Horizontal dashed lines indicate the error expected if the unperturbed (20 hour) or slowed (80 hour) growth rates were absorbed into degradation estimates. (B) Scatter plot comparing estimated degradation rates obtained from the pulse-chase (x-axis) and pulse-label (y-axis) experimental designs. Magenta line denotes y=x. (C) Scatter plots of the population mean estimated velocity (y-axis) as a function of mean abundance (x-axis) for individual genes (dots) shown separately for pulse-chase (top) and pulse-label (bottom) experiments under 4sU toxicity. (D) Vector field plots generated under 4sU slowdown for pulse-chase (top) and pulse-label (bottom) designs. Points represent individual cells, colored by the cell’s time since last division (colorbar). (E) Violin plots showing the absolute error (y-axis) between true and estimated degradation rates for genes in a mixed growth rate population (12h and 20h doubling times). Errors are stratified by gene degradation rate (x-axis) and shown for both pulse-chase (blue) and pulse-label (orange) experiments. Horizontal dashed lines mark the expected error when each subpopulation’s growth rate is absorbed into the degradation rate. (F) Scatter plots of the mean estimated velocity (y-axis) as a function of mean abundance (x-axis) for the 12-hour (green) and 20-hour (purple) doubling time subpopulations.

The two experimental designs similarly result in differences in production rate estimates. In pulse-label, *α* estimates are accurate (**Figure S2C**, bottom), though it is worth reiterating that pulse-label experiments may observe cells in a perturbed (no growth) state. For pulse-chase experiments, *γ*^∗^ is estimated from chase samples and therefore incorporates the higher growth rate, but then this value of *γ*^∗^ is applied to estimate *α* from the observed count of transcripts produced in the low-growth pulse window. Since the ground truth *α* tracks with growth rate, production rates are lower during the pulse (**Figure 2D**). Using a high growth rate *γ*^∗^ to estimate *α* from the low-growth / low-production pulse window results in systematically overestimated production rates (**Figure S2C**, top).

RNA velocity estimates from pulse-label simulations with 4sU growth defects show velocities close to zero with little apparent bias (**Figures 4C**, bottom and **S2D**, bottom). The bias is smaller than what we observed above (in the noiseless, no 4sU defect setting) due to the low growth rate (higher growth rates produce more apparent bias; **Figure 3D** vs **Figure S1F**)). In pulse-chase experiments, on the other hand, we observe a substantial negative bias in the abundance-velocity plots (**Figures 4C**, top and **S2D**, top), due to the mismatch in pulse and chase growth rates; the estimated *γ*^∗^ values incorporated a healthy 20-hour doubling time, but these values are then applied to analysis of slowed 80-hour doubling time cells, leading to systematically underestimated velocity.

Finally, trajectory plots again highlight how common estimation and interpretation procedures can lead to misleading biological conclusions. As before, ground truth velocities give trajectories proceeding from right to left in the low-dimensional visualization (**Figure 3E**, left). For pulse-label simulations with 4sU defects, cells appear to be falsely diverging along an orthogonal direction (**Figure 4D**, bottom), similar to the results above. In the pulse-chase simulation the estimated trajectories point coherently in a diagonal direction opposed to the direction of ground truth trajectories (**Figure 4D**, top), reflecting their systematically negative velocities.

### Mixed growth rate populations

We next consider populations with variable growth rates (but in the absence of 4sU defects). To do so, we generated a simulated population consisting of an equal mixture of 12-hour and 20-hour doubling time cells. Because *γ*^∗^ is estimated as a population average, the single *γ*^∗^ estimated for each gene incorporates a population average growth rate (**Figure 4E**). This is then applied to estimate *α* in single cells, each growing at a lower or higher (but never intermediate) growth rate. As a result, the estimated production rates are slightly overestimated for the slower growth population and underestimated for the faster growth population (**Figure S2E**).

These misestimated values cascade into velocity and trajectory estimation errors. The slower growth population shows a systematic negative velocity bias, while the faster growth population shows a systematic positive velocity bias (**Figure 4F**). Finally, trajectory plots from mixed growth rate simulations fail to reflect cell cycle progression, while also failing to capture differences in the two populations. Instead, cells continue to falsely appear to diverge along a direction orthogonal to cell cycle progression (**Figure S2F**).

### Effects of sampling efficiency

Above, we isolated growth rate effects by considering a noiseless scenario; we now consider more realistic settings with an error-prone experimental process. We will focus primarily on the effects of imperfect detection of transcripts produced during the pulse window, but we will also comment on the effects of low capture efficiency of transcripts overall – primarily this has the unsurprising effect of adding noise to estimated values, and we point out cases where it also introduces substantial bias (**Figure S3A,B**).

There are two commonly used approaches to detect labeled transcripts, and each can suffer from reduced efficiency. The first uses chemical conversion of a nucleotide analog (usually 4sU) that results in apparent T->C mutations in a reverse transcription dependent manner^21–23^. The second uses pulldown (usually biotin enabled) of transcripts that incorporated a modified nucleotide analog^2,52,53^. Inefficiencies, arising either molecularly or computationally, in detecting labeled transcripts in sequencing data with either method have been previously described^3,54,55^.

Here, we simulate errors with a detection efficiency parameter *δ*, where the probability that a transcript produced during the pulse window is incorrectly classified is 1 − *δ* (**Methods**). Most single cell velocity studies do not report (and potentially do not measure) this value; here we set *δ* = 0.85 as this approximate value has been reported^4,55^ and is consistent with what we observe in our studies (**Methods**).

The effects of inefficient detection differ in pulse-label and pulse-chase simulations. In chase experiments, reduced detection efficiency has little effect on *γ*^∗^ estimation (**Figures 5A,B** and **Figure S3C**); the observed abundance of labeled transcripts is reduced by the same fraction in each chase sample and the curve decays at the same rate as with perfect efficiency. Pulse-label experiments, on the other hand, calculate *γ*^∗^ from the observed decay of unlabeled transcripts during the pulse window. Since a fraction of transcripts being produced during the pulse are incorrectly classified as unlabeled, the unlabeled counts are artificially high. This fraction increases over time, resulting in a shallower decay and underestimated *γ*^∗^.

**Figure 5:**
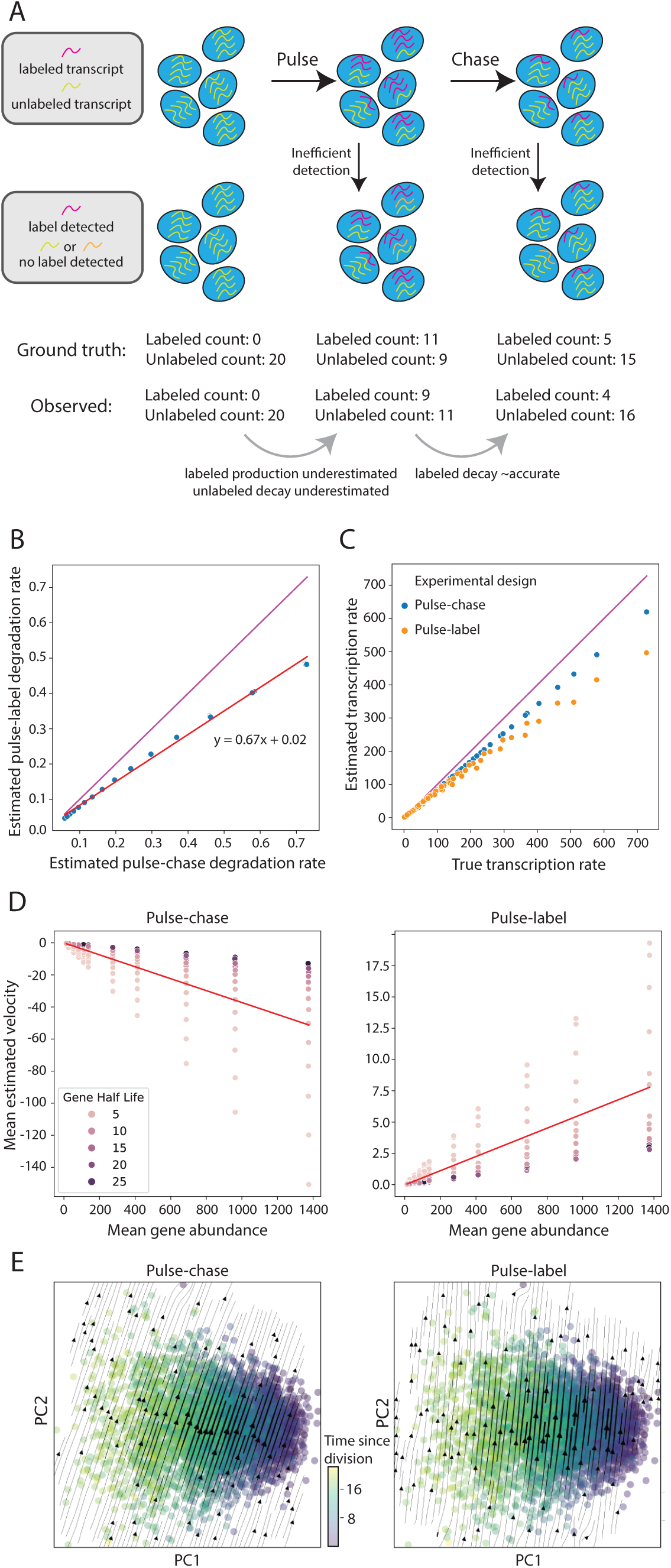
Sampling inefficiencies propagate into errors in kinetic parameter estimation. (A) Illustration of how low detection efficiency distorts the observed labeled RNA relative to the ground-truth labeling, leading to biased kinetic parameter estimates. Orange transcripts depict labeled transcripts incorrectly classified as unlabeled. (B) Scatterplot comparing degradation-rate estimates obtained from pulse-chase (x-axis) and pulse-label (y-axis) designs under low detection efficiency. Magenta line denotes y=x. (C) Estimated transcription rates (y-axis) plotted against ground truth transcription rates (x-axis) for pulse-chase (blue) and pulse-label (orange) experiments with low detection efficiency (*δ* = 0.85). Magenta line marks y=x. (D) Scatter plots of the population mean estimated velocity (y-axis) as a function of mean abundance (x-axis) for individual genes (dots), shown separately for pulse-chase (left) and pulse-label (right) experiments with low detection efficiency. (E) Vector field plots generated under inefficient labeling detection for pulse-chase (left) and pulse-label (right) designs.

For production rates, the general effect of reduced detection efficiency (or reduced capture efficiency overall) is to underestimate *α*. Production rates (and velocity) are in units of transcripts per time and are calculated from the observation of labeled transcripts produced in the pulse window. When only 85% of labeled transcripts at the end of the pulse window are detected, the estimated production rate is 85% of its true value. This is what we observe in pulse-chase experiments (**Figures 5C** and **S3D,E**). In pulse-label, *α* is further underestimated as a result of the underestimated *γ*^∗^.

The combined effects manifest in different velocity and trajectory biases, depending on experimental design. Pulse-chase experiments have a negative velocity bias, due to the underestimated production rates, and show trajectories that diverge from ground truth (**Figures 5D,E**, left and **S3F**, left). Pulse-label experiments have a positive velocity bias – both *α* and *γ*^∗^ have been underestimated, but the latter has a dominant effect – and display a prominent diagonal trajectory bias that is opposed to the direction of cell cycle progression (**Figures 5D,E**, right and **S3F**, right).

### Usage of concentrations in RNA velocity

While we have presented our analyses thus far in terms of transcript counts, in principle RNA concentrations are the more natural unit for RNA velocity since concentrations remain approximately constant across the cell cycle regardless of growth rate. We repeated our analyses but using simulated observations of mRNA concentrations and found that, despite certain improvements to RNA velocity estimation, many biases discussed above persist (**Figures 6 and S4**).

**Figure 6:**
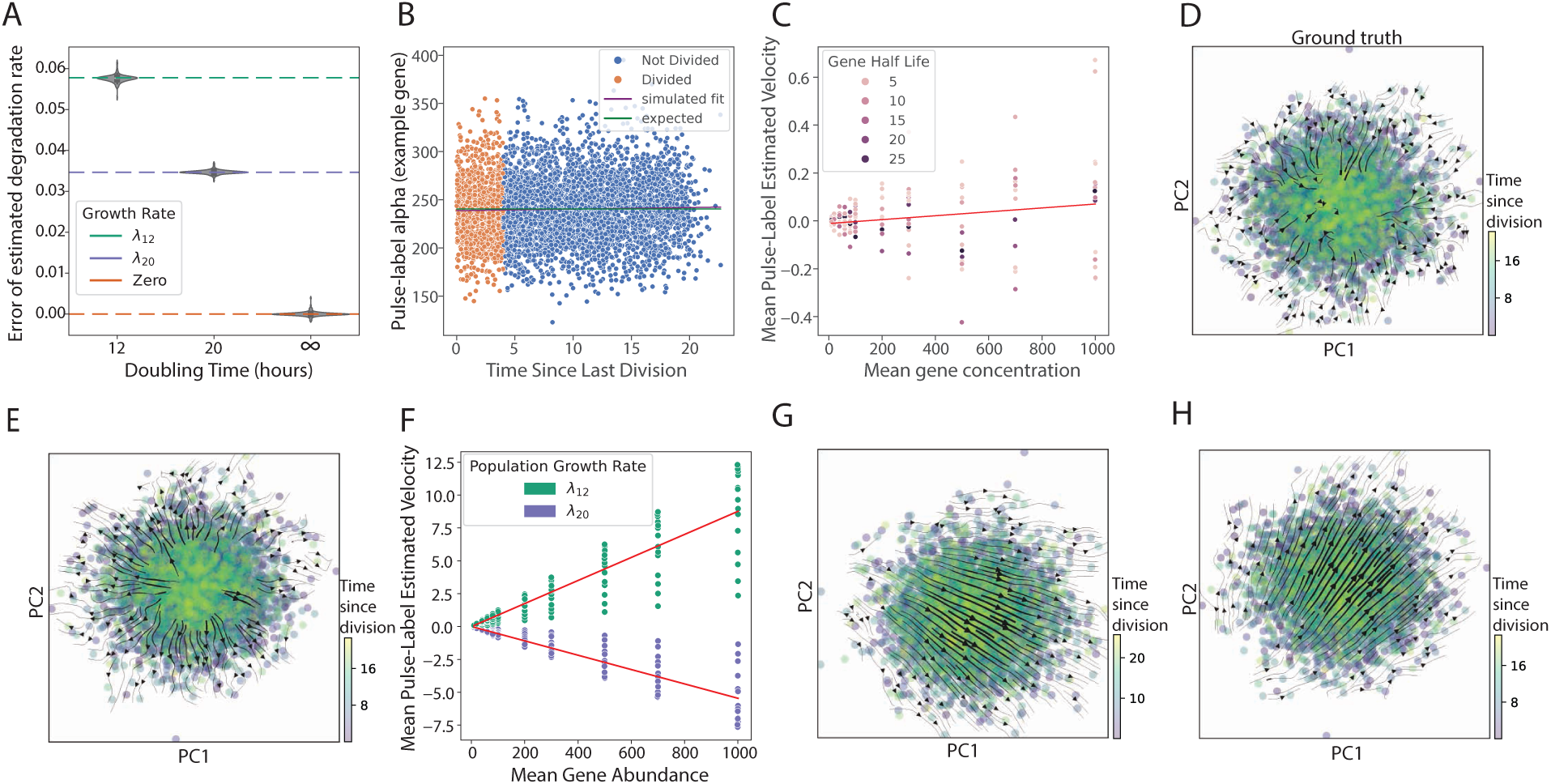
Use of concentration units in RNA velocity yields some improvements. (A) Violin plots showing the absolute error (y-axis) between true and estimated degradation rates for various growth rates (x-axis) in a pulse-label experiment with measurements in units of concentration. Horizontal dashed lines show the error expected when any one of the three growth rates is absorbed into the degradation estimates. (B) Scatter plot showing the estimated production rate (y-axis) in concentration units for a single gene (gamma=0.2057) in a population of cells (individual dots) as a function of time since last division. Curves depict a best fit to the scatter plot (purple) and the expected production rate in concentration units in exponentially growing cells. (C) Scatter plot showing the population mean RNA velocity (y-axis) as a function of mean concentration (x-axis) for individual genes (dots) in a pulse-label experiment for cells with 20-hour doubling times. (D) Vector field plots generated from ground truth velocity vectors in concentration units for each cell (dots), projected into two dimensions (PC1 and PC2). Colorbar indicates each cell’s time since last division. (E) Vector field plots generated from estimated velocity vectors in concentration units for a pulse-label experiment under baseline conditions. (F) Scatter plots of the mean estimated velocity (y-axis) as a function of mean concentration (x-axis) for the 12-hour (green) and 20-hour (purple) doubling time subpopulations in a mixed growth rate pulse-label experiment. (G) Vector field plots generated under 4sU toxicity conditions for pulse-chase experiment using concentration units. Points represent individual cells, colored by the cell’s time since last division (colorbar). (H) Vector field plots generated under inefficient labeled detection efficiency conditions for pulse-label experiment using concentration units.

Estimated degradation rates continue to absorb the cellular growth rate due to the dilution of labeled transcripts by cell volume (instead of cell division as in the case for transcript counts) (**Figures 6A and S4A**). We observed improved interpretability of RNA production rates, which scale with cell volume and thus remain constant across the cell cycle when using concentrations (**Figures 6B and S4B**). Furthermore, the consequence of incorporating the wrong growth rate due to 4sU toxicity or mixed growth rate populations, as well as the biases from imperfect labeling detection efficiency and capture efficiency, remain (**Figures 6F-H and S4E-H**). The relationship between velocity and concentration appears almost identical to that of velocity versus abundance, as the resultant velocities appear to have zero magnitude in the noiseless scenario (**Figures 6C and S4C)**, with similar distortions arising from growth rate variability and sampling efficiency (**Figures 6F and S4F-H**). This is reflected in the plot of inferred cellular trajectories. While trajectory plots inferred in the noiseless scenario from velocities estimated in units of concentration correctly capture the steady state of ground truth trajectories (**Figures 6D-E and S4D**), distortions from biases introduced by variable growth rates and sampling efficiency remain (**Figures 6G-H and S4E**).

In practice, obtaining true RNA concentrations remains a problem. The standard normalization method of scaling each cell’s UMI count to 1 million, also known as transcripts per million (TPM), reflects the relative abundance of each gene and it is tempting to think of this as a proxy for concentration. Indeed, if cells are non-proliferative and of a constant volume, TPM normalization can be reasonably thought of as converting to a proxy for concentration. In growing cells, this normalization would be effective only if the normalizing factor (total observed counts) mostly represents changes in volume rather than mostly reflecting noise.

We address this question through reanalysis of published single-cell data, and through a new experiment in which we sort and selectively sequence undivided K562 cells of increasing cell age (in cell cycle terms; **Methods**). We ask whether the normalizing factor (total counts) tends to scale appropriately with cell age (and thus size, as cells double in volume over the cell cycle; **Fig S6A**), which is a precondition if TPM normalization is to also produce a reasonable conversion to concentration. We find that TPM units are not a reliable proxy for concentrations. While several studies do exhibit an approximate doubling (using a generous “peak-to-trough” criteria), in the majority of cases total mRNA count does not double across cell cycle (**Fig S6B-I**). This result thus confirms what we believe to be the common understanding in the field: that TPM normalization primarily accounts for technical variations in total RNA per cell rather than cell volume. Below, we analyze existing metabolic labelling-based RNA velocity datasets and find that using TPM reproduces the same systematic biases seen when working with counts.

### Systematic bias evident in existing studies

A number of effects discussed above manifest in bias in abundance vs velocity plots, and so we reanalyzed data from existing studies to look for evidence of these artifacts. We analyzed four datasets (**Methods**), all with single cell RNA velocity estimates generated from nucleotide analog labeling, but using different protocols, experimental designs, and scRNA-Seq technologies^2,3,5,19^. Importantly, the cells were all in a resting state (*i.e.*, growing but not stimulated to mount a large transcriptional response, undergoing differentiation, etc.), roughly matching our simulation setting.

Our results show evidence of systematic bias in every study examined (**Figures 7A-B** and **S5**).

**Figure 7:**
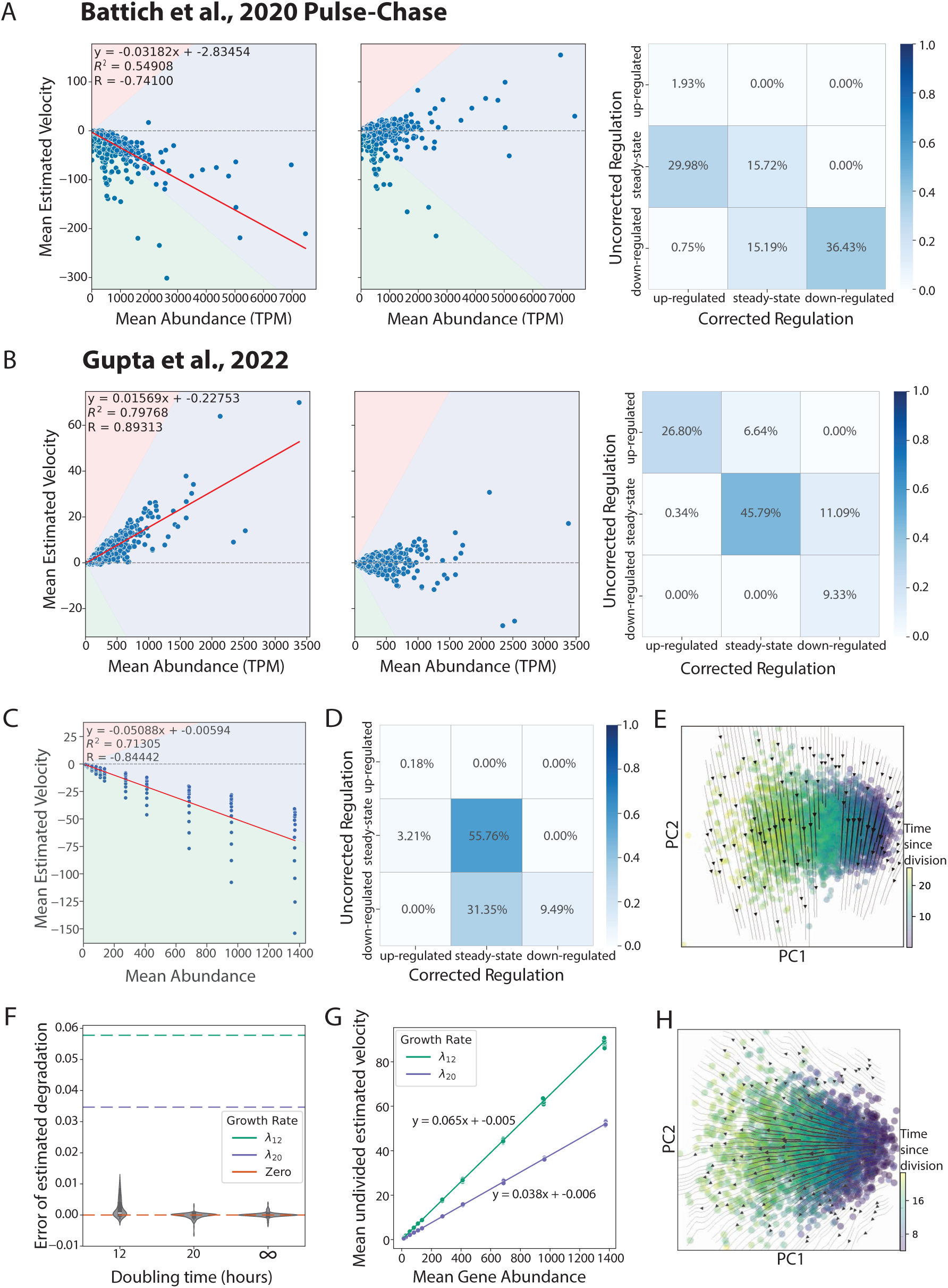
Systematic biases in existing data and possible corrections. (A and B) (left column) Scatter plots of the mean estimated velocity (y-axis) for all genes (dots) as a function of the mean gene abundance (x-axis). Red line depicts the line of best fit for this relationship. Background colors depict regulation classifications of up-regulated (orange), steady-state (blue) and down-regulated (green). (middle) Scatter plots of the mean estimated velocity corrected for systematic biases in velocity estimation (Methods) as a function of mean gene abundance. Background colors depict regulation classifications as in left plots. (right) Confusion matrix between estimated regulation classifications of single cells (y-axis) and corrected classifications after bias removal (x-axis). (C) Scatter plots of mean estimated velocity as a function of the mean gene abundance under combination 4sU toxicity and low labeled detection efficiency conditions before regression correction for cells with 20 hour doubling times. (D) Confusion matrix of estimated regulation classifications of gene-cell observations (y-axis) and corrected classifications after bias removal (x-axis). (E) Vector field plot of corrected RNA velocities in cells with combination 4sU toxicity effects and low labeled detection efficiency. Colors depict time spent in cell cycle. (F) Error plot of estimated degradation rate using undivided cells method for multiple growth rates. (G) Scatter plots of the population mean estimated velocity (y-axis) as a function of mean abundance (x-axis) for populations with 20-hour (purple) and 12-hour (green) doubling times. (H) Vector field plot of RNA velocities for cells with a 20-hour doubling time calculated using undivided cells method.

It is difficult to pinpoint the source of artifacts in each study since the confounding variables go unmeasured, but we observe a few trends. We predominantly see a negative velocity bias across studies (**Figures 7A** and **S5A**, left). Several studies also show two apparent groups of genes with a different bias in each group (**Figure S5**, left). We cannot rule out that the trends evident in abundance-velocity plots reflect “interesting” biology rather than artifacts, but our analysis nevertheless casts doubt on viability of downstream applications like trajectory mapping.

Since the sign of velocity values plays an important role in interpretation, even if quantitative values are taken with a grain of salt, we asked how often this interpretation would at least remain consistent after abundance-velocity trends are regressed out (**Figure 7A-B** and **S5**, middle; **Methods**). Setting a threshold on the magnitude of velocity to separate steady state from up- or down-regulation (**Methods**), we found the interpretation of the regulatory state of a gene changed in up to 45.9% of cases (**Figure 7A-B** and **S5**, right).

### Approaches to address growth rate bias in RNA velocity

We considered several approaches to begin addressing errors arising from the omission of growth in the velocity framework. We outline three here and elaborate on alternative future directions in **Discussion**.

To evaluate whether the computational correction described in the previous section is an accurate way to eliminate biases in the data, we regressed out the abundance-velocity trend in simulation data under a combined 4sU slowdown and detection efficiency condition. As before, RNA velocities estimated under these conditions display a clear dependence on abundance (**Figure 7C**). Regressing out the trend and classifying each gene-cell observation as steady state, up- or down-regulated, we found that 87.11% of observations are correctly identified as steady state after correction as opposed to 58.97% before correction (**Figure 7D**). However, vector field plots of corrected velocities showed that, though the spurious trend along the principal component associated with cell cycle progression (PC1) was largely eliminated, a residual trend still persists along PC2 and thus remains uncorrected (**Figure 7E**). Due to this artifact, this computational correction on its own is insufficient to correct these biases. Thus, we turn to experimental solutions.

One reason that *γ*^∗^ absorbs the growth rate is that a fixed number, rather than a fixed fraction of cells, are sampled at each time point, resulting in apparent decay of transcripts that in fact remain present in an increasingly large population of unsampled cells (**Figure 3A**). One approach, therefore, is to correct for this by either: (i) leaving the basic experimental design unchanged but measuring growth rate and multiplying the observed counts at each time point by an appropriate factor to account for the unobserved cells, or (ii) tracking the total cell population size and sampling an increasingly large number of cells (but fixed fraction) over time (potentially incurring exponentially increasing costs). In either case, the method would be subject to errors in cell counting or growth rate estimation and would not generalize to mixed growth rate populations.

An alternative approach is to experimentally isolate RNA degradation from dilution effects. In principle, this can be done by conducting a labeling experiment in which cells are sorted based on having divided or not divided. For example, in a pulse-chase experiment we could sort and sequence only the undivided population at each timepoint during the chase. Then, decay in transcript abundance should reflect true degradation and not dilution from cell division. In practice this requires also accounting for a shifting distribution of cell age in the undivided samples across time; on top of degradation processes, early chase time samples from the undivided population will have relatively more young cells (with less RNA), while late time samples will be enriched for old cells (with more RNA). We developed a method to decouple and jointly estimate degradation and growth rate, assuming parallel observations in unsorted and sorted (undivided) populations across chase time points and accounting for the shifting distribution of cell age in the latter (**Methods**).

We evaluated this approach in simulation (**Methods**), in cells with doubling times of 12 hours and of 20 hours, as above. In the noiseless setting, we were able to successfully isolate the true mRNA degradation rate from growth rate effects (**Figure 7F**). The estimated velocities are now proportional to abundance, with slope equal to the growth rate of the cell as expected (**Figure 7G**). Finally, the trajectory plots now reflect the ground truth (**Figure 3E, left**) and appropriately track cell cycle progression (**Figure 7H**). Implementing this approach in practice, however, would require an approach where counts reliably scale with cell volume, or where cellular RNA concentrations can be measured directly.

Finally, each of these corrections assumes a single growth rate across the population (**Methods**). One way to extend these solutions to mixed growth populations would be to physically or computationally sort the cells by cell type or growth rate and correct each subset independently, though we leave this to future work.

## Discussion

Single cell resolved nucleotide labeling experiments have great promise. Here, we find that existing studies show evidence of systematic bias. With the current state of the art, caution is needed and velocity estimates may not be appropriate for quantitative application in vector field mapping and downstream tasks. Our results above outline various mechanisms by which systematic biases can arise when velocity analysis is performed in growing cells. Here, we summarize our results, discuss potential solutions, and discuss the biological implications of growth rate on RNA velocity.

Our results show systematic biases in the estimation of RNA velocity kinetic parameters. First, the procedures to estimate degradation rates in single cell RNA velocity analysis produce *γ*^∗^, which cannot, in fact, be interpreted as a degradation rate. If growth rate is unmeasured and not accounted for, this value captures a mixture of distinct biological processes – RNA degradation and cell growth. If the variations in *γ*^∗^ across conditions, cell lines, or datasets simply reflected variations in RNA degradation rates, then it would be promising to study those variations to reveal principles or mechanisms of regulation, as some have proposed^56^. However, such variations can arise from differences in degradation rate, growth rate, experimental efficiency, or experimental design, complicating analysis and interpretation.

While estimated production rates can be correctly interpreted as such, variations in these estimates can similarly arise from multiple sources. Differences in single cell estimates of *α* could be due to biological phenomena at the level of the cell (growth rate or time since division) or gene (regulatory effects, presumably of primary interest), or technical effects (detection efficiency, RNA capture efficiency, or sequencing depth). Like above, with growth rate and other effects unaccounted for, it becomes difficult to cleanly study variations in production rate with an eye towards further understanding gene regulation. Moreover, it is difficult to make a case for quantitative interpretation of production rate estimates, given that total mRNA capture efficiency is low, noisy (varying from experiment to experiment or cell to cell), and typically goes unmeasured.

At the level of RNA velocity estimates, detection efficiency and growth defects have a particularly large effect of introducing systematic bias, such that the sign of velocity estimates, let alone the quantitative value, may be wrong in a large fraction of cases. Even when there is little systematic bias, it is important to interpret velocity in light of the growth rate; the same gene in two cells with the same velocity and starting abundance will see different changes in abundance if their growth rates differ. Furthermore, even when there is little systematic bias in velocity estimates, downstream analysis of trajectory plots can easily generate compelling but entirely artifactual trends.

We also make a note on the sampling procedure for RNA velocity experiments. The kinetic rate parameters of RNA velocity are generally functions of the cellular state and in practice, one should sample over short time windows over which the cell state does not change appreciably to determine these rates. It is important, however, to note that the effect of growth rate on the estimates of RNA velocity does not depend on the sampling time and the artifacts we have so far described would still be present even if short pulse or chase time windows were used. Although there may be few cell divisions or little cell volume expansion over short windows (the former relevant when working with counts, the latter when working with concentration), one ends up dividing the observed decrease in abundance (or concentration) by a small window of time, exactly reproducing the same error when long windows are used. Cell state transitions that occur during the experimental window would potentially add compounding errors on top of the artifacts we describe, but they are not the source of these artifacts.

While our simulation framework provided a controlled environment to identify growth rate effects on RNA velocity, we performed certain abstractions of the underlying biology for practical reasons. Specifically, we modeled cells with singular doubling times, whereas *in vivo*, cells may exhibit large cell-cell heterogeneity in growth rate. However, the use of singular growth rates allowed for clearer illustration of growth rate effects. In addition, simulated cells were set to divide on time-based mechanisms of duration doubling time plus some variation. However, cells can exhibit volume homeostasis mechanisms rather than the time-based mechanisms as in our simulation^57–59^. As individual cell doubling times had relatively small variance, we believe this has minimal impact on the simulation. Additionally, we modeled cell division with even transcription partition whereas, *in vivo*, partitioning during cell division is stochastic and often asymmetric^60^. However, as a general case, modeling symmetric cell division simplifies the simulation with relatively little drawback. Finally, we did not simulate the effects of gene-copy number changes. However we do not believe that it plays a significant role in mammalian cells as the process of gene replication only takes a small portion of the cell cycle, thus leaving “gene allocation fractions” unchanged before and after replication^43^. Nevertheless, we believe these abstractions of the underlying biology do not alter our central findings.

Above, we discussed two options to explicitly account for growth in experimental design (computational and experimental correction). These options are particularly limited as they do not allow for heterogeneity in growth rate. Here, we outline further possibilities for addressing core problems. In particular, changing the experimental modality to *in situ*, imaging-based measurements could facilitate several important alterations. There is no method yet capable of measuring velocity *in situ*, but important steps have been taken, critically showing that labeling based transcriptional readouts can be made *in situ*^61^. Further development to allow imaging-based velocity measurements would allow (i) coupling with live tracking^62^ to identify and (computationally) isolate cells that did not divide during the pulse window, in place of the sorting and sequencing approach above; (ii) coupling transcriptional information with measurement of cell volume, allowing for computation of expression levels and kinetic rates in units of concentration; (iii) higher fidelity measurements – FISH-based spatial transcriptomics can capture a substantially higher fraction of mRNA molecules than single cell sequencing, reducing the effect of low capture rates and detection efficiency.

The choice of pulse-label or pulse-chase experiment can also play a big role in the sign and magnitude of bias. The two designs consistently give different estimates of the same parameters. In several cases the pulse-label design appeared preferable. The impacts of 4sU growth defects (**Figure 4C**) and label detection efficiency (**Figure 5D**) were not as large in pulse-label as in pulse-chase designs. However, pulse-label experiments were not without bias, and it is worth reiterating that pulsing with a nucleotide analog can cause growth defects, meaning cells studied in the pulse window can be in a perturbed, no growth state.

In studies of heterogenous populations, the mixture of growth rates causes unique effects because the population average growth rate is absorbed into degradation estimates. With two populations, this leads to systematically biased estimates in opposite directions for each cell in the two groups. With current, scRNA-Seq based velocity methods, there is no easy fix. Ad-hoc solutions, like estimating the growth rate of each cell type, then using scRNA counts to identify cell type and correct for type-specific growth rate, are possible. In our view, this is ultimately unsatisfying but may nevertheless be better than completely ignoring growth effects.

Further, explicit representation of growth rate suggests an interesting biological mechanism of gene regulation: growth rate can be a global regulator of gene inducibility. We see this by examining the differential equation from the mathematical framework for RNA dynamics in growing cells (**Methods**). In slow growing cells a change in affinity Δ*a* can result in a large shift in abundance, while the same induced Δ*a* results in a small shift in abundance in fast growing cells (**Figure S7A**). We also test this in simulation by simulating genes with the same starting abundance and same induction, then examining how the new steady state abundance varies by growth rate (**Figure S7B, Methods**). We found that in both cases, the same degree of induction or repression (i.e., the absolute change in production rate) will result in different changes in abundance, depending on the growth rate, with the effect being particularly larger for long half-life genes (**Figure S7C-D, Methods**). We note that this phenomenon has also recently been observed in *E. coli*^63^.

Though single cell RNA velocity methods have found utility in many applications, there remains substantial skepticism regarding their quantitative utility. Our hope is that the results presented here spur interest in development of refined methods that can ultimately bring the promise of RNA velocity to fruition.

## METHODS

### Review of existing literature

To contextualize the 43 publications that develop, benchmark, or review RNA velocity methods, we queried OpenAlex (openalex.org) on March 19, 2026 and identified 4,836 unique publications that have cited at least one of these 43 studies. Next, to investigate how many of these papers are susceptible to growth rate effects, we restricted our analysis to the 200 most-cited publications published after 2021, 171 of which performed velocity analysis (the remainder being reviews or tangential citations). Finally, we used Claude (Opus 4.8; accessed March 19, 2026) to classify these publications into four categories based on biological system studied: Proliferating, mixed, post-mitotic only, and not applicable. Of the 171 papers that applied RNA velocity to a biological system, 167 (97.7%) involve proliferating cells either exclusively or as part of a mixed population.

### Modeling steady state expression with cell growth

Our model of RNA kinetics in growing cells is largely the same framework developed by Lin and Amir^42^. In particular, we follow the setting where ribosomes are limiting for growth and RNA polymerase is limiting for mRNA production, though we note the other scenarios outlined by Lin and Amir will generally produce the same phenomena outlined in Results. Below we describe our implementation of this model, along with modifications to introduce bursty transcription.

To model steady-state transcriptional dynamics, we simulated mRNA abundance trajectories for each gene under a kinetic framework defined by its instantaneous transcription rate, *α*(*t*), a constant population degradation rate, *γ*, and the current mRNA abundance, *x_i_*. The temporal change in abundance follows

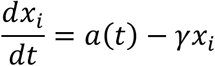

In this formulation, transcription is constrained by the finite number of RNA polymerase (RNAP) molecules available to each cell, with RNAP abundance increasing proportionally with cell growth over the cell cycle. This coupling between RNAP availability and cell volume enables the model to capture the coordinated increase in mRNA production observed during cell growth. Promoter activity is additionally modeled using a two-state bursting process, providing a more realistic representation of transcriptional kinetics. Details of the RNAP scaling scheme, two-state kinetics, and parameter selection are provided below.

### Modeling cell volume dynamics

Because transcription is limited by the number of RNAP, and because RNAP abundance scales with cell volume, we begin by modeling how cell volume changes over the cell cycle. Changes in volume, *V*(*t*), are proportional to the rate of population growth and the current volume, yielding an exponential curve. The growth rate, *λ*, is set by the cell doubling time, *T_d_*, yielding a 2-fold increase in volume over the cell cycle:

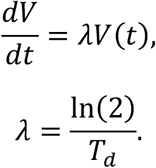

### Defining gene kinetic parameters

We next specify gene-specific kinetic parameter.

In the model described by Lin and Amir^42^, the transcription rate, *α*(*t*), of a gene is:

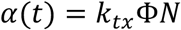

Where *k_tx_* is the (constant) rate of transcription of a single RNAP, *N is* the total number of RNAPs (proportional to cell volume), and Φ is the “gene allocation fraction”, or the constant coarse-grained quantity that models what fraction of the limited RNAP pool will be allocated to a given gene.

Firstly, *N*, the total number of RNAPs, is proportional to cell volume, *V*(*t*):

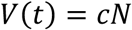

Where *c* is a constant.

In our simulations, *k_tx_* Φ, and *c* are constant, and so we implicitly absorb these values into a new constant describing the initial transcription rate, *α*_0_.

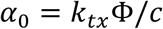

Thus, we define *α*_0_ explicitly by assigning each gene an initial abundance, *A*_0_, and a half-life, *t*_1/2_. Degradation rates, *γ*, for each gene were computed as 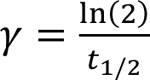. Combined with the cell growth rate, *λ*, these values determine a gene-specific initial transcription rate *α*_0_ = *A*_0_(*γ* + *λ*). At any time *t*, the effective transcription rate scales with cell volume:

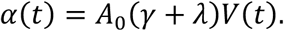

We selected cells with doubling times, *T_d_*, of 12 hours (*λ* = ∼0.0578 ℎ*r*^−1^) and 20 hours (*λ* = ∼0.0347 ℎ*r*^−1^). In addition, informed by half-lives calculated from Schofield et al., we selected 15 half-lives, *t*_1/2_, that were uniformly spaced on a log scale between 1 and 30 hours^22^. In addition, we selected a range of initial abundances informed by existing literature, *A*_0_ = {10,20,40,60,80,100,200,300,500,700,1000}^64^. From this, we defined 165 genes with every combination of initial abundance and half-life.

### Simulation of bursty RNA kinetics

To incorporate transcriptional bursting kinetics, we used a two-state promoter model in which each gene alternates between an inactive (OFF) and active (ON) state. In this model, transcripts are only produced during ON periods at synthesis rate, *k_syn_*. Promoter switching is governed by two rate constants: the activation rate, *k_on_*(*t*), which determines how frequently a promoter enters the ON state, and the inactivation rate, *k_off_*, which determines how quickly it returns to the OFF state. The expected dwell times in each state are therefore 1/*k_on_*(*t*) (time spent OFF), and 1/*k_off_* (time spent ON).

While in the ON state, the promoter produces transcripts at synthesis rate, *k_syn_*. The burst size, *b*, is defined as the expected number of transcripts produced during a single ON period:

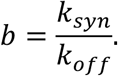

Thus, the mean transcription rate at time *t* results from the combined contributions of the average burst size, *b*, the average duration the gene spends in the ON state per cycle, 1/*k_off_*, and the average duration the gene spends in the OFF state per cycle, 1/*k_on_*(*t*):

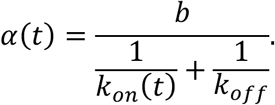

To ensure that the bursting process is fully consistent with the gene-specific transcriptional program generated by cell growth (via *V*(*t*)), we solved this relationship for

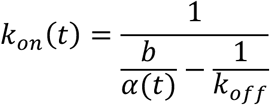

This guarantees that, for each gene at every time point in the simulation, the promoter switching rates produce the exact time-dependent transcription rate required by the model. Thus, promoter activity adapts dynamically as the cell grows and transcriptional capacity increases with volume.

### Selection of bursting parameters

We defined the burst size as a fixed fraction of the gene’s initial abundance:

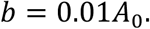

This choice allowed for the scaling of bursts with gene expression level, as larger-abundance genes produce proportionally larger bursts. In addition, setting *b* to 1% of the initial abundance leads to numerically stable results.

We defined *k_off_* such that the burst duration would be brief relative to the mRNA lifetime:

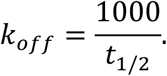

Thus, we set a burst duration of 1/*k_off_* = *t*_1/2_/1000. In addition, this allows for *k_on_* to be well defined, as *k_off_* must be high enough such that the denominator of 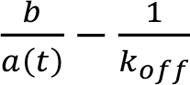 remains nonzero and positive for all simulated values.

### Running the simulations

#### Simulation of cells at steady state

We simulated this model using the stochastic simulation algorithm (SSA)^65^. We simulated 70 lineages, each with varying initial start times from [0, *T_d_*], and for every cell we tracked all 165 simulated genes. The simulation was then run for eight times the doubling time of the cell. Every gene in the origin cell was set to its *A*_0_ in the OFF state before the simulation began.

At every time step, we simulated three molecular processes. First, the 2-state switching mechanism in which genes transition between OFF and ON states. If a gene is in the OFF state, then it has propensity *k_on_*(*t*) of switching to the ON state. This process is time dependent as time-dependent changes in overall transcription rate, *α*(*t*), propagate into *k_on_*(*t*), as described earlier. If the gene is in the ON state, then it has propensity *k_off_* of switching to the OFF state. Second, we simulate transcriptional bursts. When a gene is in the ON state, it has propensity *k_syn_* of producing one new transcript. Finally, we simulate transcript degradation. At all times, we model transcript degradation as a first order reaction with propensity *γx*, where *x* is the current number of transcripts.

We simulated metabolic labeling by assigning each newly synthesized transcript to either a labeled or unlabeled pool, depending on the current simulation time. During the defined labeling period, *k_syn_* generated labeled transcripts. Otherwise, newly synthesized transcripts were assigned to the unlabeled pool.

Additionally, we simulated a growing population through cell division by triggering a cell division event after *T_d_* + *ε* hours, in which *T_d_* is the cell doubling time and *ε* is normally distributed noise with variance 0.005*T_d_*. During division, both cell volume *V*(*t*) and transcript counts *x* were partitioned evenly between the daughter cells, preserving labeled and unlabeled identities.

#### Simulation of transcript labeling experiments

We simulated two types of labeling experiments, pulse-label and pulse-chase, using experimental designs analogous to prior work. For pulse-label experiments, cells were exposed to a 4-hour pulse, during which all newly transcribed mRNA were assigned to a labeled population, and pre-existing transcripts remained unlabeled. Cells were sampled hourly throughout the pulse.

For pulse-chase experiments, a 4-hour pulse was followed by a 20-hour chase. During the chase, newly transcribed mRNA was assigned to the unlabeled population, and the labeled mRNA produced during the pulse was allowed to decay. Cells were sampled hourly throughout the chase.

To model incomplete labeling detection, we introduced a parameter, *δ*, which represents the probability that a labeled transcript is correctly identified as labeled during sequencing. During our simulation of the sequencing process, each labeled molecule captured was assigned a 1 − *δ* probability of being misclassified as unlabeled.

To model the effects of sparse sequencing, we introduced a capture-efficiency parameter,

*π*, which represents the probability that a given transcript is detected by the sequencing technology. After simulating the true transcript counts for each gene-cell pair, we applied Bernoulli thinning in which each transcript was retained with probability *π* and discarded otherwise. This produced sparse count matrices that reflect reduced molecular capture efficiency.

#### 4sU slowdown effect

To model the growth-inhibitory effects of the commonly used nucleotide analog 4sU, we introduced a slowdown factor, *τ_s_*, which proportionally increases the cell’s doubling time. The slowed growth rate, *λ_s_*, was therefore applied only during the 4sU pulse period by defining a time-dependent growth:

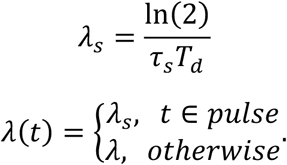

Cell volumes were then calculated under the new time-dependent growth rate, simulating slowed increases in volumetric expansion and RNAP availability during the pulse.

Because division timing depends on the instantaneous growth rate, we adjusted each cell’s time-to-division based on the fraction of the cell’s nominal (un-slowed) cycle spent in the slowed state, *f_s_*. Thus, the cell’s effective doubling time was updated:

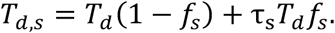

Given the slowed growth rate and volume trajectory, we recalculated transcription rates ensuring that the transcriptional output decreased coherently alongside growth:

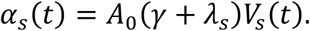

This reduced transcription rate yielded a correspondingly reduced burst frequency, *k_on_*_,*s*_(*t*), computed by substituting *α_s_*(*t*) into the bursting model.

Together, these modifications allowed 4sU exposure to slow cellular growth, delay division timing, and proportionally reduce transcriptional activity during the labeling period.

#### Simulations of mixed growth populations

To model heterogeneous populations with distinct growth rates, we simulated two independent cell populations that differed only in their intrinsic growth rates. Specifically, one population was assigned a doubling time of 12 hours (or growth rate of *λ*_12_) and the other a doubling time of 20 hours (or growth rate of *λ*_20_), with all other parameters held constant across groups. Both populations contained the same set of genes, defined by their initial abundance, *A*_0_, and half-life, *t*_1/2_. Each population was simulated separately through the full transcriptional and labeling framework described above.

Following simulation, the two populations were combined into a single population to generate a mixed growth rate population.

#### Simulations of Non-Growing Cells

To examine how RNA velocity behaves in the absence of cell growth, we simulated a non-growing condition in which the growth rate was set to zero (*λ* = 0). In this case, cell divisions were disabled, and the cell volume remained constant, *V*(*t*) = 1. Because transcription in our model scales with *γ* + *λ* and cell volume, the transcription rate simplifies to a constant value.

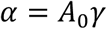

Given this constant transcription rate, we calculate a constant burst propensity, *k_on_*, using the relationship defined previously. All other kinetic parameters are identical to those used in the growing-cell simulations. Experimental labeling designs and all downstream velocity analyses were implemented in the same manner as for the growing conditions, allowing direct comparison between growth conditions.

#### Simulations of RNA concentrations

To examine if RNA velocity measurements would improve with the use of concentration units rather than RNA counts, we calculated RNA concentrations for a cell, *x_conc_*, using simulated RNA counts, *x*, and simulated cell volume, *V*.

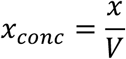

Given these gene concentrations, we performed RNA velocity analyses to arrive at estimated production and degradation rates, as well as estimated RNA velocities.

### Analysis of simulations

#### RNA velocity estimation

Based on the observed labeled or unlabeled counts for each gene, we estimated a population-level degradation rate *γ*^∗^ by fitting an exponential decay model to decaying transcript counts over time. For pulse-label experiments, we computed the mean unlabeled abundance, *u_av_*_g_(*t*), across cells at each sampled time point, *t*, during the pulse. The degradation rate, *γ*^∗^, was then obtained by nonlinear least-squares regression of the following model to the series of measured time-points:

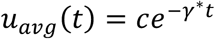

Similarly, for pulse-chase experiments, we computed the mean labeled abundance, *l_av_*_g_(*t*), across cells at each sampled time point, *t*, during the chase. We then performed nonlinear least-squares regression of the following model to obtain the degradation rate:

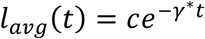

Given the estimated population average degradation rate, *γ*^∗^, we estimate single-cell transcription rates using the observed number of labeled transcripts present at the end of a pulse labeling period of duration *t_p_*. For both pulse-label and pulse-chase assays, this corresponds to the labeled counts at the end of each pulse, which we denote as *l*(*t_p_*). Then, the transcription rate estimate is given by:

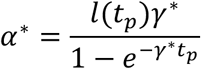

Finally, we obtained single-cell velocity vectors by applying the inferred gene-level degradation rate, *γ*^∗^, together with each cell’s estimated transcription rate, *α*^∗^, and observed total abundance, *x*. For each gene-cell pair, (*i*, *j*), velocities, *v*, are computed as follows, yielding a velocity vector for each cell:

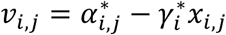

In order to investigate how growth rate influences RNA velocity estimates, we calculated the average abundance and average estimated velocity for each gene across all cells:

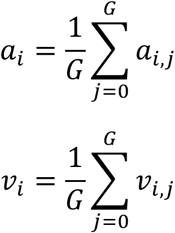

We assessed whether growth-rate differences introduced systemic biases into the velocity estimates by plotting gene-wise mean abundances against gene-wise mean velocities.

#### Vector Field Plots

We embedded the simulated cells using principal component analysis (PCA) of the observed count matrix (cells by genes, independent of label status), retaining the first two principal components for visualization. Estimated RNA velocity vectors for each cell were projected into this space using the PC loadings, and the resulting vector field was visualized using the velocity_embedding_stream function from scVelo 0.3.3^7^.

To generate ground-truth vector field plots, we identified each cell’s simulated state 0.5 hours into the future and excluded any cells that divided within that interval. Ground truth velocity vectors were computed as the difference between the future and current state for each cell. These ground-truth velocity vectors were projected into PC space using the same loadings and visualized identically to the estimated velocities.

### Simulations of Undivided Cells

To determine if isolating undivided cells can isolate true degradation rates from growth rates, we simulated this experimental condition and analyzed its impact on RNA velocity estimates.

#### Estimation of degradation rates

Late-stage cells possess higher total RNA counts, and, therefore, a higher number of labeled transcripts. Because these cells exit the tracked population earlier during the chase due to division, the remaining population becomes increasingly skewed toward cells that began the chase with fewer labeled counts. This sampling bias artificially inflates the estimated degradation rate. In order to resolve this bias, we performed the following correction.

#### Modeling sampling bias

The rate of change of labeled counts, *l*, during a pulse of duration *t_p_* follows:

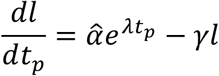

Where *α*a = *α*_0_*e^λa^*, or the transcription rate for a given cell of age *a* at the start of the pulse. With initial condition, *l*(0) = 0, we model labeled counts during the pulse with:

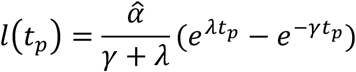

Thus, the number of labeled counts in a cell, *l*(*t_p_*), is proportional to the transcription rate, *α*a. Here, *α*a has a distribution that follows from the age distribution of cells at the beginning of the pulse window. The density distribution, *f*, of ages, *θ*, follows:

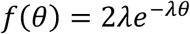

As time progresses during the chase, the population of undivided cells will have a shifting distribution, *f*(*θ*). At later chase times, the remaining undivided population will bias more towards cells that were early in their cell cycle at the start of the pulse and will thus have lower values of *α* and fewer labeled counts.

The expectation of synthesis rates across undivided cells at chase time *t_c_* is calculated by integrating over the surviving initial age range, [0, *t_d_* − *t_c_*]:

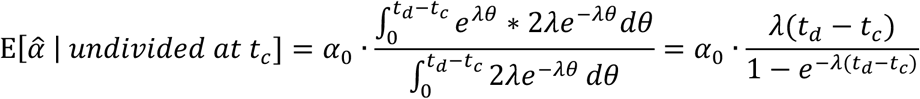

Because *l*(*t_p_*) ∝ *α*a, the labeled counts in the chase inherit this relationship, scaled by mRNA decay, *e*^-*γtc*^:

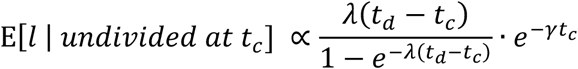

Finally, substituting *t* = ln(2)/λ and simplifying, we arrive at:

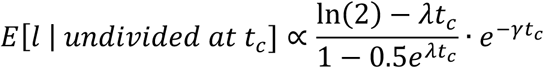

#### Isolating degradation rates

In order to isolate degradation rates from growth rate using this correction, we begin by first calculating decay of mRNA transcripts in the total (undivided and divided) population, *γ^*^_tot_*. As in our previous findings, *γ*^∗^ = *γ* + *λ*. Solving for true degradation rate, we arrive at:

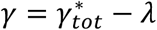

Substituting this into the previous equation yields the fitting equation:

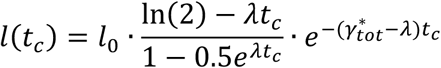

Finally, we estimate the remaining global parameter, *λ*, by fitting the equation above, and then we recover the true degradation rates, *γ*, per gene via *γ* = *γ*^∗^ − *λ*.

#### Estimation of Transcription Rates

We begin with the rate of change of labeled counts during the pulse:

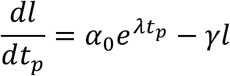

With initial condition, *l*(0) = 0

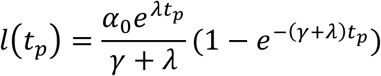

Finally, we can calculate estimated transcription rate, *α*^∗^, using the combined degradation rate and growth rate term, *γ* + *λ*:

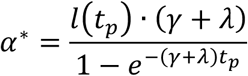

#### Estimation of RNA Velocities

In order to calculate RNA velocities with isolated degradation rates, we used the calculated true degradation rate, *γ*, and the estimated production rate, *α*^∗^:

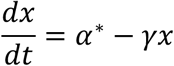

### Analysis of existing datasets

#### Studies examined

In order to investigate whether growth rate effects are present in existing studies, we reanalyzed four publicly available datasets: Battich et al. (2020, GSE128365)^2^, which includes both pulse-label and pulse-chase experiments; Liu et al. (2023, GSE197667)^5^; Gupta et al. (2022, GSE202292)^19^; and Qiu et al. (2020, GSE141851)^3^. All datasets were processed using a standardized velocity pipeline as described below.

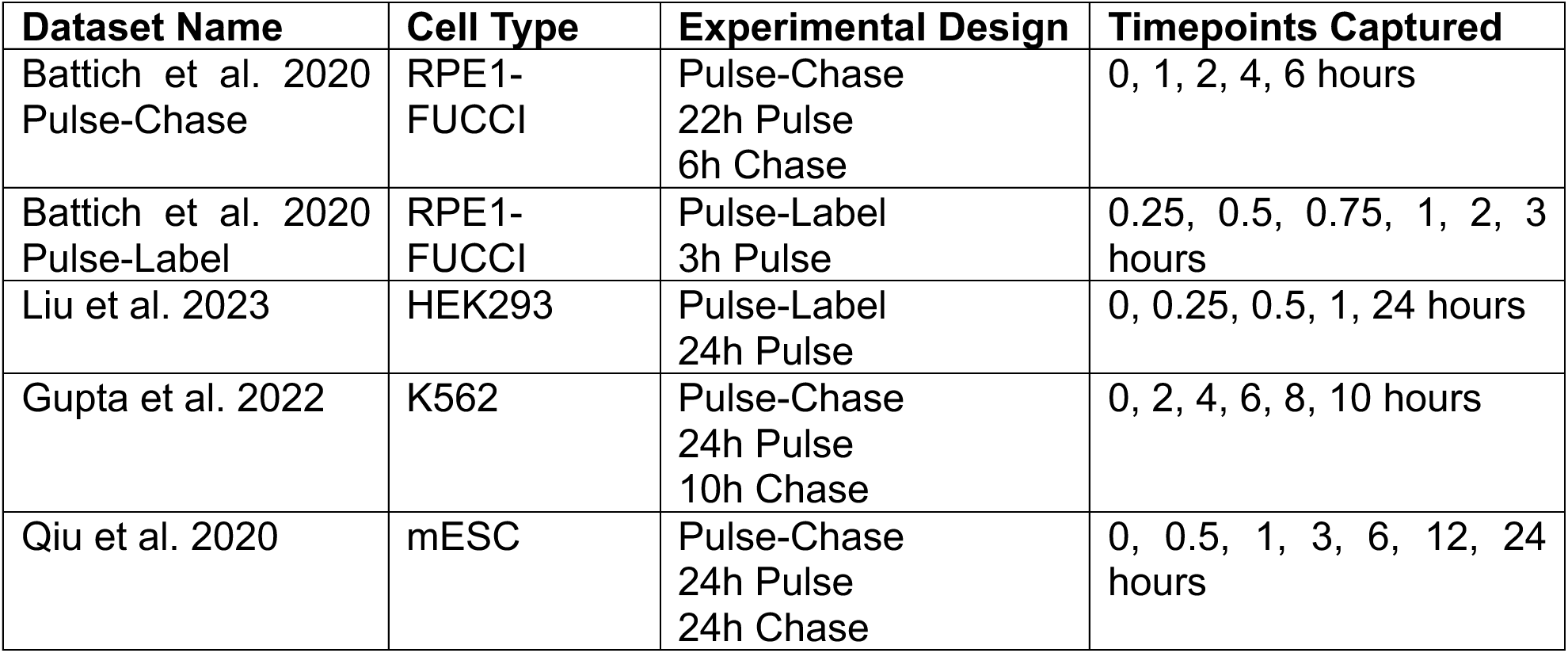

#### Filtering and normalization

We removed sparsely detected genes and low-quality cells by retaining only genes expressed in at least 100 cells, and cells expressing at least 500 genes. We “transcripts per million” (TPM) normalized each cell to its total transcript count. Let *X_i,j_* denote the count for gene *i* in cell *j*, and let *X*_>_ = ∑*^G^ X_i,j_* be the total count across all *G* genes for each cell. The TPM value for each gene was then defined as:

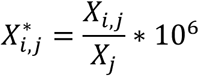

This is intended to correct for variations in both cell size and mRNA capture efficiency. The normalized matrices were subsequently used for RNA velocity estimation and downstream analyses.

#### RNA velocity analysis

We applied the same velocity estimation procedure used for simulated data to each experimental dataset. Briefly, for each gene we estimated a population average degradation rate by fitting an exponential decay model to the transcript population that decreases over time in each experimental design. We used unlabeled transcripts for the pulse-label datasets, Battich et al (2020) Pulse-Label and Liu et al (2023), and labeled transcripts for the pulse-chase datasets, Battich et al (2020) Pulse-Chase, Gupta et al. (2022), and Qiu et al (2020). Transcription rates were then computed from the labeled counts at the end of the labeling period. Finally, RNA velocity estimates were calculated using the calculated transcription and degradation rates, as well as the total mRNA counts in the cell.

Dataset-specific considerations were addressed as needed. In the Battich et al. (2020) Pulse-Label dataset, the earliest 15-minute pulse sample was excluded due to incomplete 4sU incorporation. Mitochondrial genes were filtered out of datasets, as they exhibited a uniquely strong negative trend.

#### Bias Correction Analysis

To correct for systematic dependencies between gene abundance and estimated velocities, we modeled the mean velocity of each gene, *v_i_*, as a linear function of its mean abundance, *x*o*_ι_*. We fit a regression of the form

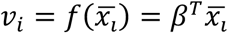

This regression captures the global abundance-velocity trend across genes. We then corrected single-cell velocities by subtracting the expected velocity predicted by *f*. For each gene *i* in cell *j*, the residual of the cell-specific velocity from its abundance describes the corrected velocity, *v_i,j,corrected_*.

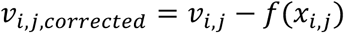

Several datasets exhibited multiple distinct abundance-velocity trends (three trends in Liu et al. 2023 and two in the Battich et al. 2020 pulse-label dataset). In these cases, genes were manually separated into trend groups based on the visual structure of the abundance-velocity relationship, and regressions were fit independently within each group to obtain cluster-specific corrected velocities.

We next assigned each gene-cell observation to a regulatory category using the combination of an absolute velocity threshold, *ε*, and a relative threshold, *ε*_%_, which was scaled by mean abundance. The absolute threshold prevents low-abundance genes from being classified as up- or down-regulated due to noise, while the relative threshold ensures that higher abundance genes require proportionally larger velocity deviations to be classified as non-steady state. We calculate a relative velocity, *r_i,j_*, which is used to define the regulation state of a given gene, *i*, in a given cell, *j*:

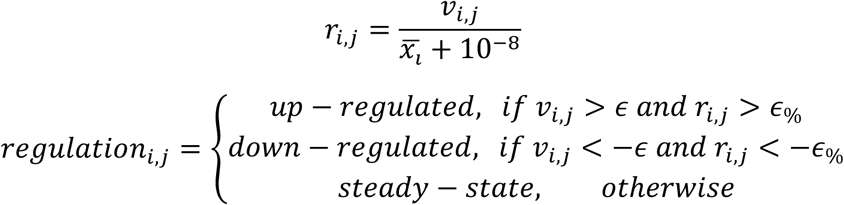

The same classification scheme was applied to corrected velocities, *v_i,j,corrected_*. Observations with zero total transcript counts were excluded from classification.

Using these assignments, we quantified the impact of the bias correction by computing the fraction of gene-cell observations whose regulatory assignment changed after correction, and by constructing a confusion matrix comparing pre- and post-correction regulations.

### Gene induction experiments

#### Induction model

To model transcription induction in growing cells, we introduced the perturbation Δ*α*_0_, which increases the initial transcription rate of a gene:

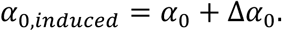

To investigate how this induction affects the new steady-state abundance of a gene (at the beginning of the cell cycle), we define the initial transcription rate of a gene without intervention:

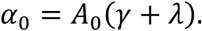

The induced transcription rate, *α*_0,*induced*_, can be rewritten as:

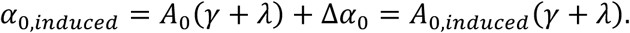

Solving for the new steady state abundance, *A*_0,*induced*_, gives:

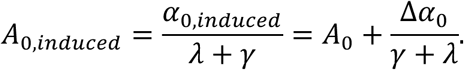

Thus, induction increases abundance by an amount determined not only by the induction strength Δ*α*_0_, but also by the effective loss rate *γ* + *λ*. Because faster growing cells dilute transcripts more rapidly, higher growth rates lead to smaller increases in steady-state abundance for the same transcriptional perturbation.

#### Induction simulations

To explore this result in simulation, we performed induction experiments for all genes under three growth conditions: doubling times of 12-hours and 20-hours, and a no-growth condition in which *λ* = 0 and cell volume remains constant. For each condition, we applied a constant increase in the initial transcription rate, Δ*α* = 7.5, to every gene in every cell. Induction was introduced 50 simulation hours before final sampling, which ensures that all genes reach their new steady-state abundances. Due to the slower effective transcript turnover in no-growth cells (where the loss rate is *γ* rather than *γ* + *λ*), induction was applied 150 simulation hours before final sampling to allow these cells sufficient time to reach steady state.

Because transcription rates directly affect the bursting model, this induction also resulted in recalculation of the burst propensity parameter *k_on_*(*t*) according to the previously defined relationship. All other kinetic parameters were held constant during induction.

After the 50-hour settling period, cells were sampled, and the new steady-state abundances of induced genes were compared across the three growth conditions. In order to enable fair comparison between growing cells and non-growing cells, changes in abundance for growing cells were normalized by cell volume.

### Cell Culture

K562 (CCL-243 ATCC) myelogenous leukemia cells were purchased from ATCC. Unless otherwise stated, K562 cells were cultured in RPMI-1640 (Cytiva, SH30255.FS) supplemented with 10% fetal bovine serum (Millipore Sigma, F0926) and 1% penicillin-streptomycin (Sigma-Aldrich, P4333) and grown at 37°C and 5% CO2.

### CFSE Proliferation Assay

K562 cells were stained with CellTrace CFSE (Thermo Fisher Scientific, C34554) per manufacturer’s instructions and resuspended to a concentration of ∼0.2 million cells/ml in RPMI-1640 (no glutamine, Thermo Fisher Scientific, 21870092) supplemented with 4mM L-glutamine (Cytiva, SH30034.01), 10% fetal bovine serum, and 1% pencillin-streptomycin. For 4-thiouridine (4sU) and uridine-treated cells, 4sU (Sigma-Aldrich, T4509) and uridine (Sigma-Aldrich, U3003) were added to separate cell cultures to a final concentration of 100μM and 10mM respectively. Cells were cultured continuously for 90h, with cell density maintained at 0.4 million cells/ml by adding fresh media every 20-24h. Untreated, 4sU-treated, and uridine-treated cells were sampled 10min (used as positive control) and 90h after staining and incubation at 37°C and fixed in 10% formalin (Fisher Scientific, BP531) in PBS. The fluorescence intensity of the cells was measured by flow cytometry using the Attune NxT flow cytometer.

### 24h Pulse-label Experiment

K562 cells were grown until ∼0.5 million cells/ml and pulsed with 100μM 4sU for 24h, with regular media change every 4h. 2 replicates were performed for this pulse-label experiment. A control sample pulsed with DMSO (instead of 4sU) was included to control for exposure to DMSO (which 4sU is dissolved in) and to measure the basal rate of T-to-C mutations in non-4sU-treated cells. Cells were harvested at the end of 24h, washed once with cold HBSS (Thermo Fisher Scientific, 14025134) and fixed in 80% methanol (Millipore Sigma, 494437) at −20°C for 30min. Chemical conversion of 4sU was performed by adding 20μl of 0.5M iodoacetamide (Thermo Fisher Scientific, A39271) to 1ml of cell suspension each, incubating overnight at room temperature with gentle agitation. The reaction was quenched by resuspending the cells in quenching buffer (DPBS (Thermo Fisher Scientific, 14190250), 0.1%BSA (Sigma-Aldrich, A6003), 1U/μl RNase inhibitor (Thermo Fisher Scientific, AM2696), 100mM DTT (Sigma-Aldrich, D0632)) for 5min at room temperature. The cells were then resuspended in resuspension buffer (DPBS, 0.01%BSA, 0.5U/μl RNase inhibitor, 1mM DTT) and passed through a cell strainer before continuing with the preparation of 10x Chromium Next GEM Single Cell 3ʹ v3.1 sequencing libraries according to standard protocol. Libraries were sequenced with a sequencing configuration of 28:10:10:90 cycles for Read1:i7:i5:Read2.

### Pulse-Chase Experiment with Undivided Cells

K562 cells were grown until ∼0.5 million cells/ml and pulsed with 100μM 4sU for 4h. A control sample pulsed with DMSO was included as in the pulse-label experiment. At the end of the pulse, cells were washed twice with HBSS and labelled with CellTrace CFSE as per manufacturer’s instructions. Cells were then resuspended in media supplemented with 10mM uridine and chased for another 24h. Cells were sampled at 0h (10min after staining with CFSE), 4h, 8h, 12h, and 30h, stained with LIVE/DEAD fixable violet dead cell stain (Thermo Fisher Scientific, L34964), and fixed in 80% methanol at −20°C for 30min before transferring to −80°C for longer storage (<1 week). Chemical conversion of 4sU was performed as in the pulse-label experiment, quenched, and then resuspended in resuspension buffer before proceeding to cell sorting using a Sony SH800S cell sorter.

Since K562 has an estimated doubling time of ∼16-18h, it can be assumed that most cells have undergone at least 1 round of cell division during the chase in the 30h sample. The density of cells was also verified by cell counting to have approximately doubled from the 0h to the 30h timepoint. To isolate the population of undivided cells from the 0h, 4h, 8h, and 12h chase samples, cells were gated on the CFSE fluorescence channel to only include cells with fluorescence greater than the top ∼1% of the 30h sample. 500k cells were isolated from each sample, with flow cytometry data captured for the first 100k cells. Both sorted and unsorted populations of chase samples for the 0h, 4h, 8h, and 12h timepoints were then used in the preparation of 10x Chromium GEM-X Single Cell 3’ v4 sequencing libraries according to standard protocol. Libraries were sequenced using a NovaSeq X with a sequencing configuration of 28:10:10:150 cycles for Read1:i7:i5:Read2.

### Processing of Sequencing Data

Raw FASTQ files were preprocessed and mapped to human reference genome GRCh38.p14 using STARsolo v2.7.10b^66^ with HI, AS, and MD tags and –soloFeatures Gene GeneFull Veloctyo included in addition to default parameters to obtain spliced, unspliced, and ambiguous count matrices. Quantification of labeled and unlabeled reads was performed with dynast v1.0.1(https://github.com/aristoteleo/dynast-release) with GENCODE annotation human release 48, generating cell barcode by gene matrices of total, labeled, and unlabeled UMI counts contained in a single .h5ad file for each sample.

### Estimating Detection Efficiency

Since the median RNA half-life has been reported to be around ∼1-2h^22^, most RNA would be labeled after a 24h pulse with 4sU. The detection efficiency of the 24h pulse-label experiment was estimated by taking the mean across all cells of labeled UMI counts divided by total UMI counts for all genes with half-lives previously confidently estimated (r2>0.6) to be between 1-4h. The detection efficiency was calculated to be between 0.82-0.85.

### Estimating Cell Size during Pulse-Chase Experiment

As cells grow and divide across the chase timepoints from 0-12h, the undivided population of cells should be enriched in cells at later stages of the cell cycle at later chase timepoints.

Assuming the cell as a perfect sphere, the forward scatter signal of the sorted cells is proportionate to the cross-sectional area of the cell^67^. The volume of the cells was calculated from the FSC-A channel according to the formula:

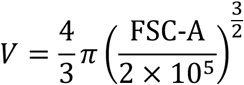

In an asynchronous population of cells, the density of cells along the cell cycle phase is exponentially distributed:

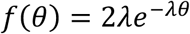

Where *θ* ∈ [0, *T*) is the time from cell division, 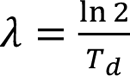 is the growth rate of the cell and *T_d_* is the doubling time of the cell. The volume *x* of the cell at *θ* is:

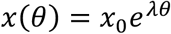

Where *x*_0_ is the initial volume of the cell at *θ* = 0. In the sorted population of cells, divided cells are removed from the population. The normalized density of the remaining undivided cells at time t is given by:

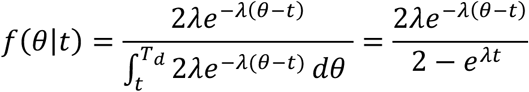

Where *θ* ∈ [*t*, *T_d_*). The expected cell volume is thus:

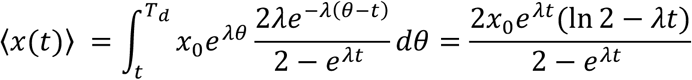

Where 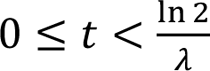.

The doubling time of the cells was calculated by fitting the mean values of the cell sizes across the sorted chase timepoints from 0-12h to the equation of the expected cell volume.

### Total RNA Counts during Pulse-Chase Experiment

As the number of reads per cell increases, the number of new unique molecules sequenced per cell decreases due to sequencing saturation. As such, at high read depth, the number of UMIs detected per cell does not scale linearly with the actual total RNA per cell. To account for sequencing saturation, an estimation of total UMI counts per cell was calculated by using the following formula:

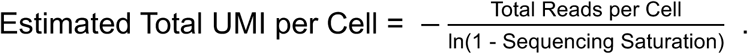

### Comparison of Total RNA against Cell Cycle Phase or Volume

To investigate how total UMI counts per cell changes with cell cycle phase or volume, we compared data obtained from this experiment with single-cell sequencing data from Gupta et al., 2022 K562 cells^19^, Battich et al., 2020 RPE1-FUCCI cells (pulse-only)^2^, 10x Genomics GEM-X v4 dataset HEK293T/NIH3T3 cells^68^, Aissa et al., 2021 PC9 day 0 cells^69^, Lederer et al., 2024 RPE1-FUCCI cells^70^, and Capolupo et al., 2022 human fibroblast cells^71^. The data from this experiment, as well as data from Gupta et al., 2022 and Battich et al., 2020 were processed as described above. The original raw data from 10x Genomics GEM-X v4 Single Cell 3’ dataset was aligned to human reference genome GRCh38.p14 and mouse reference genome GRCm39 and processed using STARsolo v2.7.10b with –soloFeatures Gene GeneFull Velocyto to obtain spliced, unspliced, and ambiguous count matrices. Since the 10x Genomics GEM-X v4 dataset contains a 1:1 mix of human HEK293T cells and mouse NIH3T3 cells, cells were split into human or mouse cells if they contained >0.9 human genes or <0.1 human genes respectively. Processed data with spliced and unspliced count matrices for Aissa et al., 2021, Capolupo et al., 2022, and Lederer et al., 2024 datasets were obtained from the VeloCycle study^72^ and are available on Zenodo at https://doi.org/10.5281/zenodo.12517650. VeloCycle was used to assign cells to a continuous cell cycle phase φ, using genes from the ‘large’ gene set available in ‘velocycle.utils’, after filtering for genes with >0.1 mean unspliced UMIs per cell or with >0.3 mean spliced UMIs per cell, with 5000 training steps used for manifold learning. Fold change of total UMI counts were calculated from the peak/trough ratio from binned means of total UMI counts per cell. All datasets were split into 48 bins except for Capolupo et al., 2022 (20 bins) and Battich et al., 2020 (47 bins) datasets to ensure ∼60 cells per bin.

## DATA AND CODE AVAILABILITY

Code used for simulations and existing data analyses is available at https://github.com/algo-bio-lab/velocity-with-growth.

Raw datasets from public sources (Battich et al. (2020, GSE128365)^2^; Liu et al. (2023, GSE197667)^5^; Gupta et al. (2022, GSE202292)^19^; Qiu et al. (2020, GSE141851)^3^, and 10x Genomics GEM-X v4 Single Cell 3’ 1:1 Mixture of Human HEK293T and Mouse NIH3T3 Cells^68^) are publicly accessible through their respective repositories. Processed data for PC9 day 0 cells^69^, human fibroblast cells^71^, and RPE1-FUCCI cells^70^ are available on the Zenodo repository for VeloCycle^72^ at https://doi.org/10.5281/zenodo.12517650.

## ACKNOWLEDGEMENTS

We thank members of the Cleary lab for fruitful discussions regarding this manuscript. This work was supported by NIH grant R35GM160123.

## SUPPLEMENTAL FIGURES

**Figure S1:**
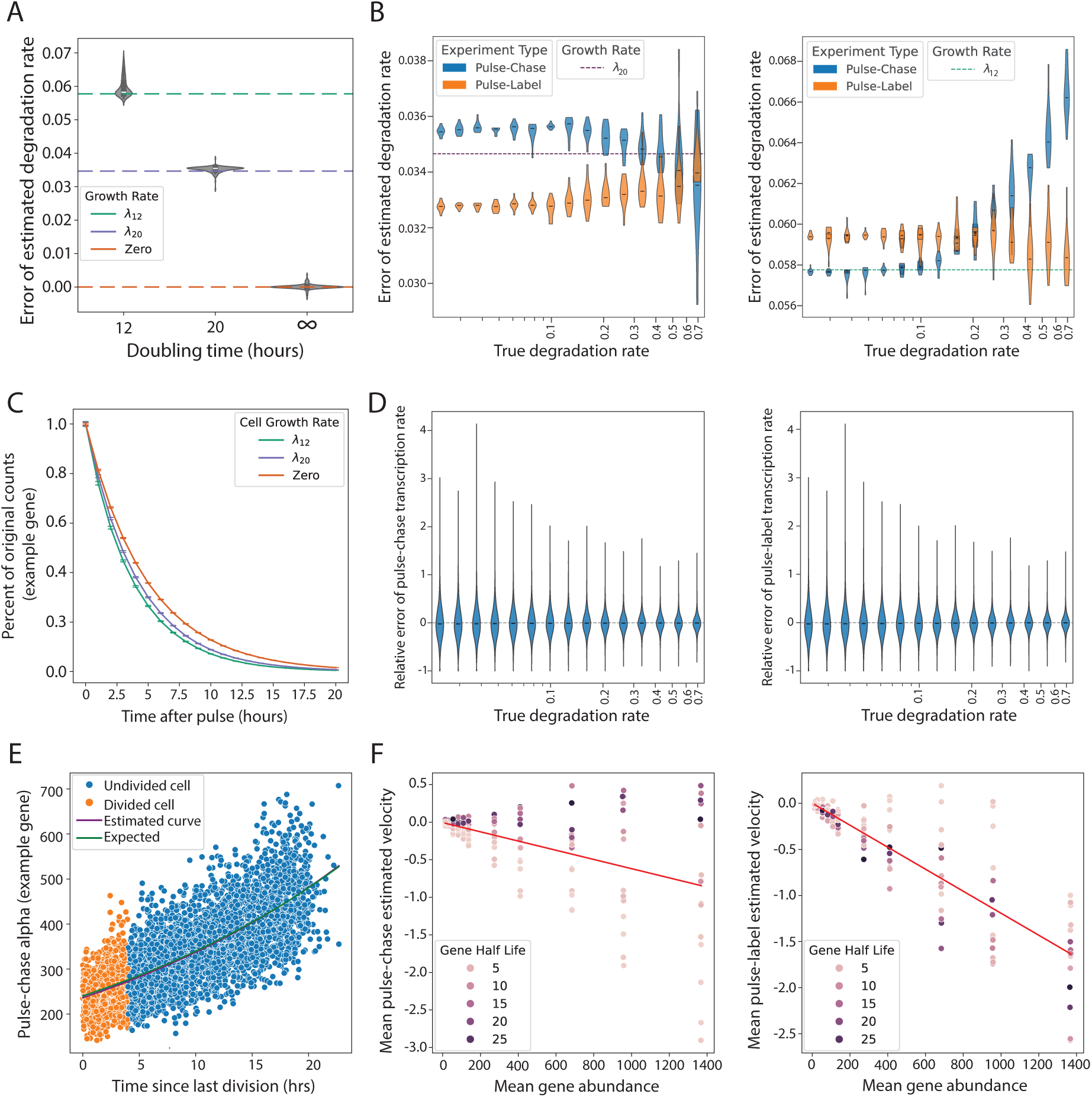
Robustness of growth rate incorporation into degradation rate (A) Violin plots showing the absolute error (y-axis) between true and estimated degradation rates for non-growing cells or with a 12- or 20-hour doubling time in a pulse-chase experiment. Horizontal dashed lines show the error expected when any one of the three growth rates is absorbed into degradation estimates. (B) Violin plots showing the absolute error (y-axis) between true and estimated degradation rates across genes for cells with a 20-hour doubling time (left) and a 12-hour doubling time (right). Errors are stratified by ground-truth degradation rates (x-axis) and shown for both pulse-chase (blue) and pulse-label (orange) experiments. Horizontal dashed lines indicate the error expected if the growth rate was absorbed into degradation estimates. (C) Decay curves depicting observed degradation of labeled counts for an example gene (*γ*=0.2057) over time after pulse (x-axis). (D) Violin plots showing the relative error of estimated transcription rates in pulse-chase (left) and pulse-label (right) experiments, stratified by ground truth degradation rates. (E) Scatter plot showing the estimated production rate (y-axis) for a single gene in a population of cells (individual dots) as a function of each cell’s time since last division (x-axis). Colors for each dot indicate whether a cell underwent division between the start of the chase and the time of sampling. Curves depict a best fit to the scatter plot (purple) and the expected production rate in exponentially growing cells (green). (F) Scatter plots of the population mean estimated velocity (y-axis) as a function of mean abundance (x-axis) for individual genes (dots) in pulse-chase (left) or pulse-label (right) experiments for cells with 12-hour doubling times.

**Figure S2:**
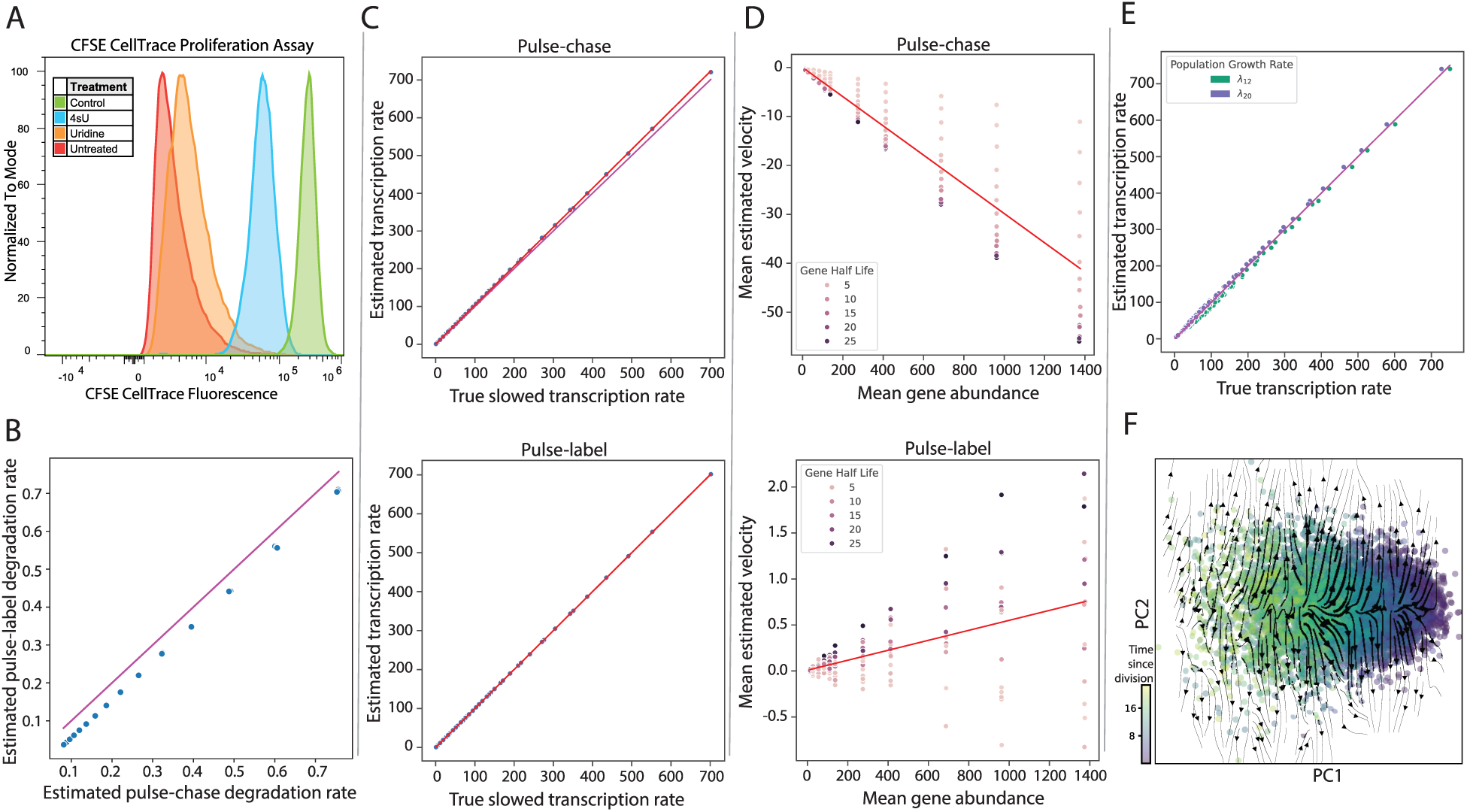
Additional analyses of variable growth rates effects on velocity estimation (A) Relative fluorescence intensity of CFSE CellTrace-stained K562 cells treated with 100µM 4sU, 10mM uridine, or without treatment, grown continuously for 90h. Control population reflects freshly stained (undivided) cells. (B) Scatter plot comparing estimated degradation rates obtained from the pulse-chase (x-axis) and pulse-label (y-axis) experimental designs (for 12-hour growth rate cells). Magenta line denotes y=x. (C) Estimated transcription rates (y-axis) plotted against ground truth transcription rates (x-axis) for pulse-chase (top) and pulse-label (bottom) experiments under a 4sU-induced slowdown. Magenta line marks y=x. (D) Scatter plots of the population mean estimated velocity (y-axis) as a function of mean abundance (x-axis) for individual genes (dots) shown separately for pulse-chase (top) and pulse-label (bottom) experiments for cells with 12-hour doubling times under 4sU toxicity. (E) Estimated transcription rates (y-axis) plotted against ground truth transcription rates (x-axis) for a mixed growth rate population. Subpopulations of cells with 12-hour (green) and 20-hour (magenta) doubling times are separated. (F) Vector field plots generated for mixed growth-rate populations from a pulse-chase experiment.

**Figure S3:**
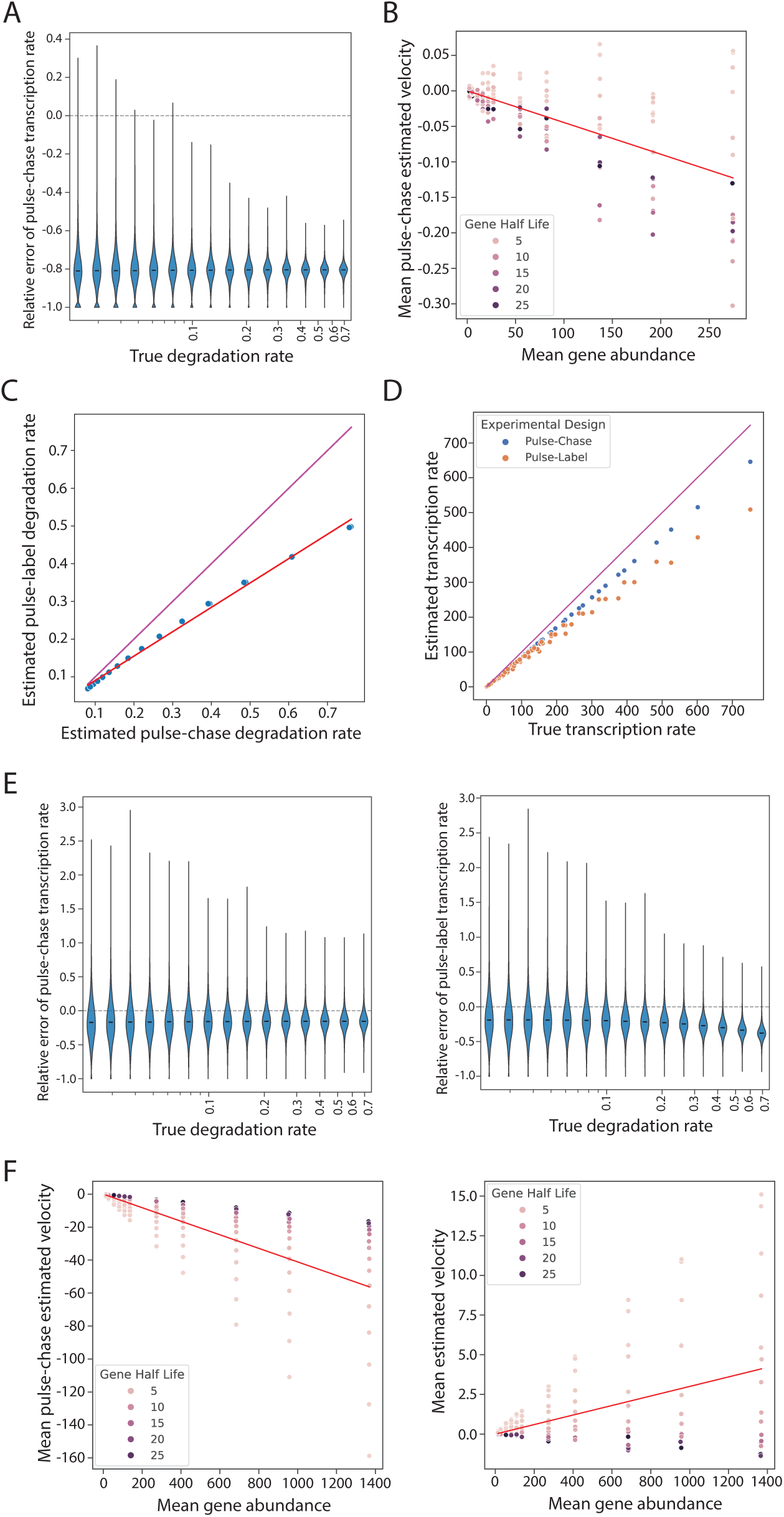
Additional analyses of sampling effects on velocity estimation (A) Violin plots showing the relative error (y-axis) between true and estimated transcription rates across degradation rates (x-axis) and (B) scatterplots of the mean estimated velocity (x-axis) as a function of mean abundance (y-axis) under 20% capture efficiency in a pulse-chase experiment. (C) Scatterplot comparing estimated degradation rates from pulse-chase (x-axis) and pulse-label (y-axis) experiments under low labeling detection efficiency in cells with a 12-hour doubling time. Magenta line denotes y=x. (D) Estimated transcription rates (y-axis) plotted against ground truth transcription rates (x-axis) for pulse-chase (blue) and pulse-label (orange) experiments in cells with a 12-hour doubling time with low detection efficiency. (E) Violin plots showing the relative error (y-axis) between true and estimated transcription rates across degradation rates (x-axis) under low labeling detection efficiency for pulse-chase (left) and pulse-label (right) experiments in 20-hour doubling time cells. (F) Scatter plots of the population mean estimated velocity (y-axis) as a function of mean abundance (x-axis) for individual genes (dots) in cells with a 12-hour doubling time, shown separately for pulse-chase (left) and pulse-label (right) experiments with low detection efficiency.

**Figure S4:**
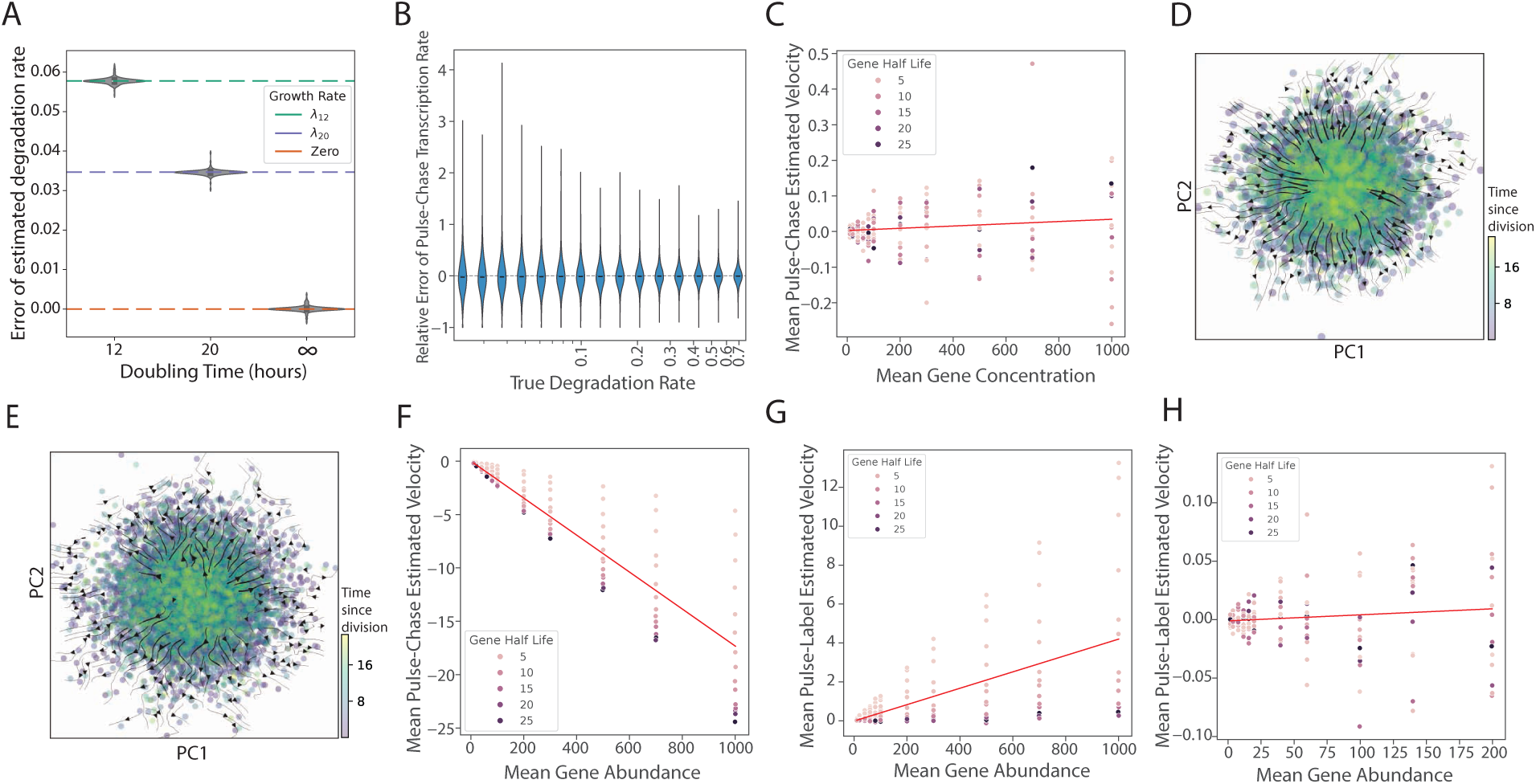
Additional analyses of usage of concentration units in RNA velocity. (A) Violin plots showing the absolute error (y-axis) between true and estimated degradation rates for various growth rates (x-axis) in a pulse-chase experiment. Horizontal dashed lines show the error expected when any one of the three growth rates is absorbed into the degradation estimates. (B) Violin plots showing the relative error (y-axis) of estimated transcription rates in pulse-label experiment using concentration units, stratified by ground truth degradation rates (C) Scatter plot showing the population mean RNA velocity (y-axis) as a function of mean concentration (x-axis) for individual genes (dots) in a pulse-chase experiment for cells with 20-hour doubling times. (D) Vector field plots generated from estimated velocity vectors in concentration units for a pulse-chase experiment under baseline conditions. Points represent individual cells, colored by the cell’s time since last division (colorbar). (E) Vector field plots generated under mixed growth rate condition conditions for pulse-label experiment using concentration units. (F) Scatter plots of the mean estimated velocity (y-axis) as a function of mean concentration (x-axis) under 4sU toxicity conditions for a pulse-chase experiment. (G) Scatter plots of the mean estimated velocity (y-axis) as a function of mean concentration (x-axis) under inefficient labeled detection conditions for a pulse-label experiment. (H) Scatter plots of the mean estimated velocity (y-axis) as a function of mean concentration (x-axis) under low capture efficiency conditions for a pulse-label experiment.

**Figure S5.**
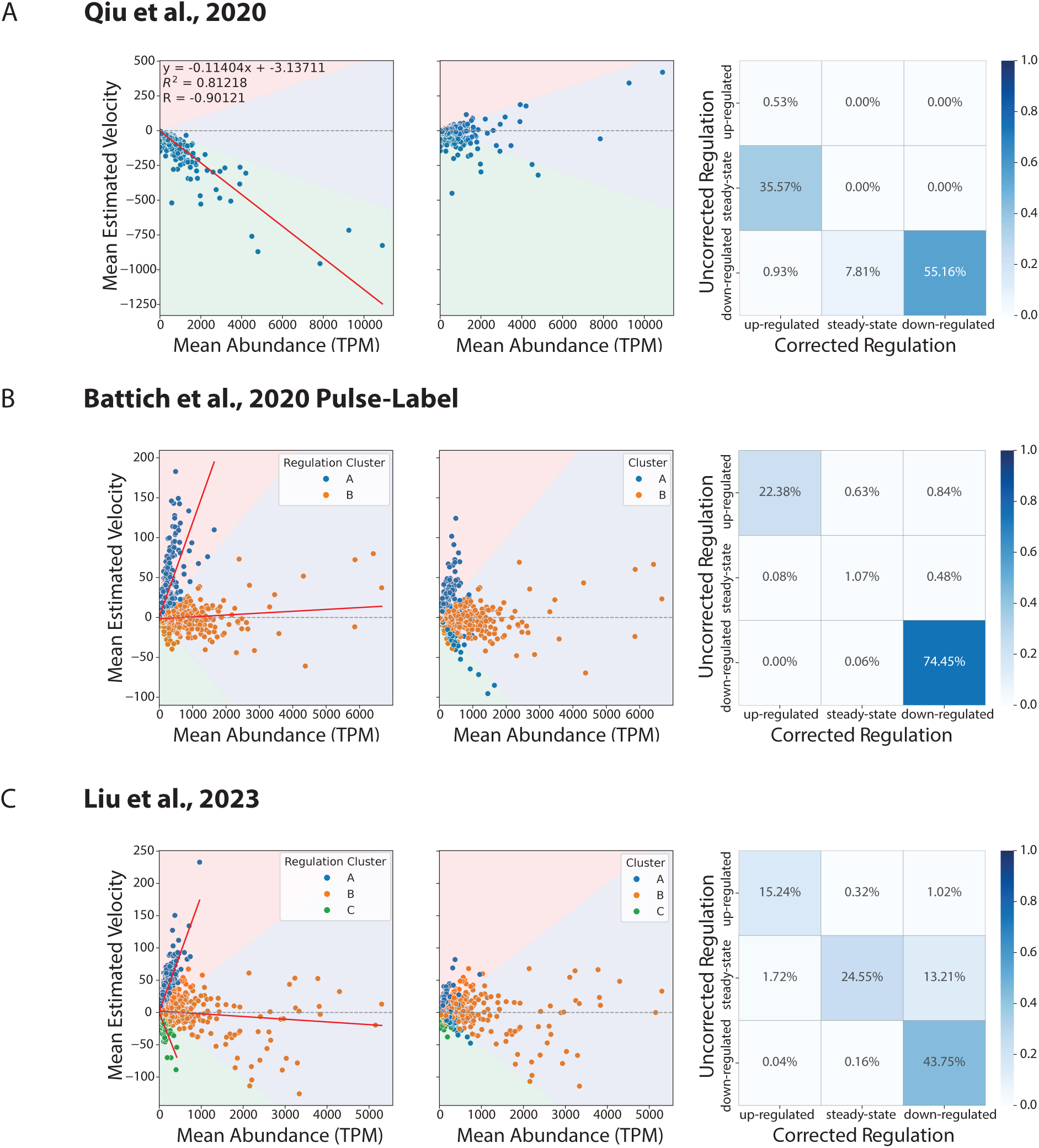
Different groups of systematic biases in existing data of RNA velocity estimation. (left) Scatter plots of the mean estimated velocity (y-axis) for all genes (dots) as a function of the mean gene abundance (x-axis). Red line depicts the line of best fit for this relationship. Background colors depict regulation classifications of up-regulated (orange), steady-state (blue) and down-regulated (green). (middle) Scatter plots of the mean estimated velocity corrected for systematic biases in velocity estimation (Methods) as a function of mean gene abundance. Background colors depict regulation classifications as in left plots. (right) Confusion matrix between estimated regulation classifications of single cells (y-axis) and corrected classifications after bias removal (x-axis)

**Figure S6:**
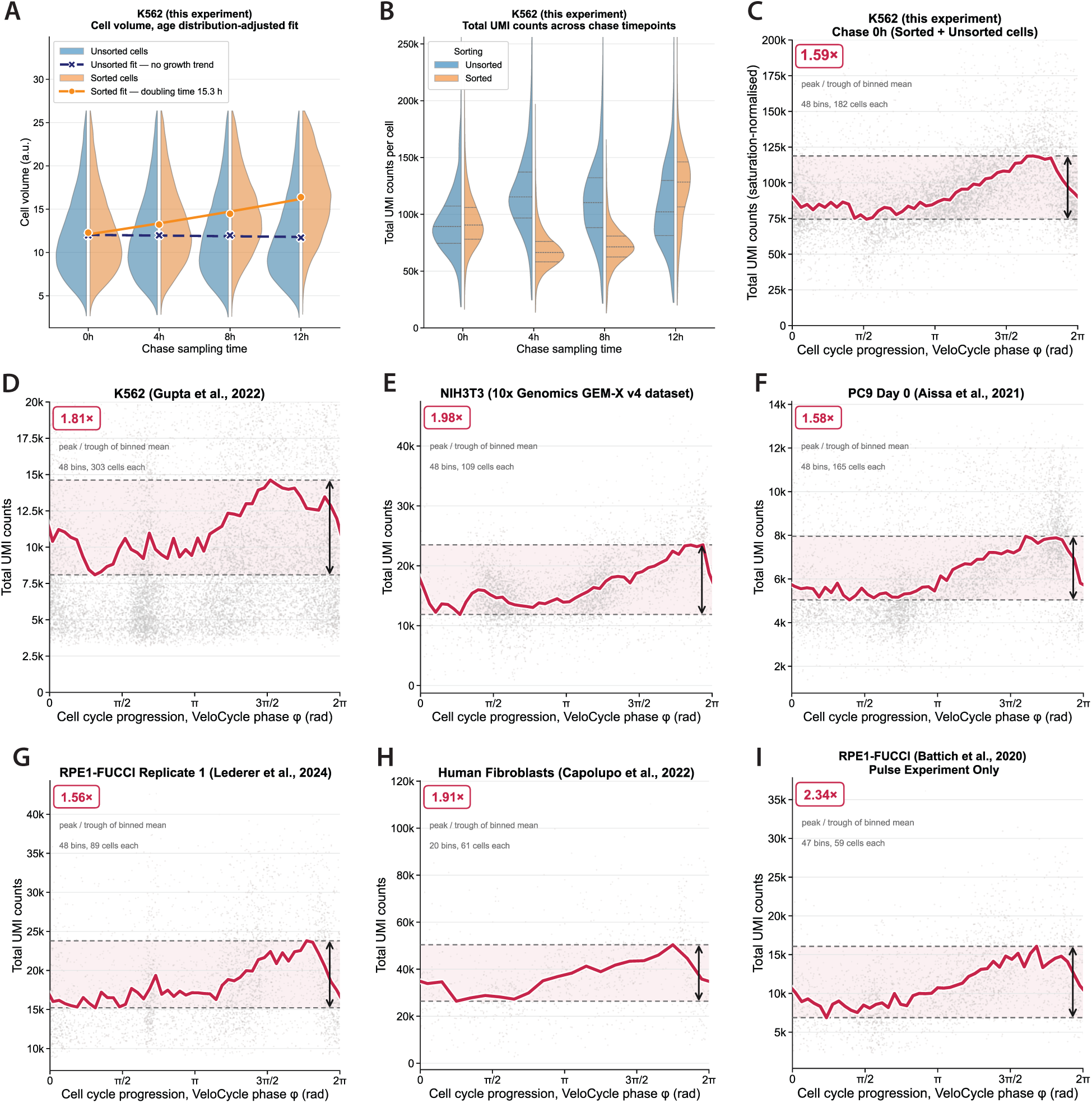
scRNA-Seq UMI Counts do not double with cell size. (A) Relative cell volumes of unsorted and sorted (undivided) K562 cells across 4sU chase timepoints. Age distribution-adjusted exponential fit of mean cell sizes across chase timepoints shows an estimated doubling time of 15.3h. (B) Total UMI counts per cell distributions, normalized for sequencing saturation, across unsorted and sorted (undivided) K562 cells across 4sU chase timepoints. Total RNA counts of sorted populations do not show a uniform increase in UMI with cell size. (C) Total UMI counts of unsorted and sorted (undivided) K562 cells from this experiment at 0h of chase timepoint, normalized for sequencing saturation, across cell cycle coordinates estimated with Velocycle. (D) Total UMI counts of K562 cells from Gupta et al., 2022 dataset, across cell cycle coordinates estimated with Velocycle. (E) Total UMI counts of NIH3T3 cells from 10x Genomics 10k 1:1 Mixture of Human HEK293T and Mouse NIH3T3 Cells GEM-X v4 2024 dataset, across cell cycle coordinates estimated with Velocycle (F) Total UMI counts of PC9 Day 0 cells from Aissa et al., 2021 dataset, across cell cycle coordinates estimated with Velocycle (G) Total UMI counts of RPE1-FUCCI cells (replicate 1) from Lederer et al., 2024 dataset, across cell cycle coordinates estimated with Velocycle (H) Total UMI counts of human fibroblast cells from Capolupo et al., 2022 dataset, across cell cycle coordinates estimated with Velocycle (I) Total UMI counts of RPE1-FUCCI cells from Battich et al., 2020 dataset, across cell cycle coordinates estimated with Velocycle

**Figure S7:**
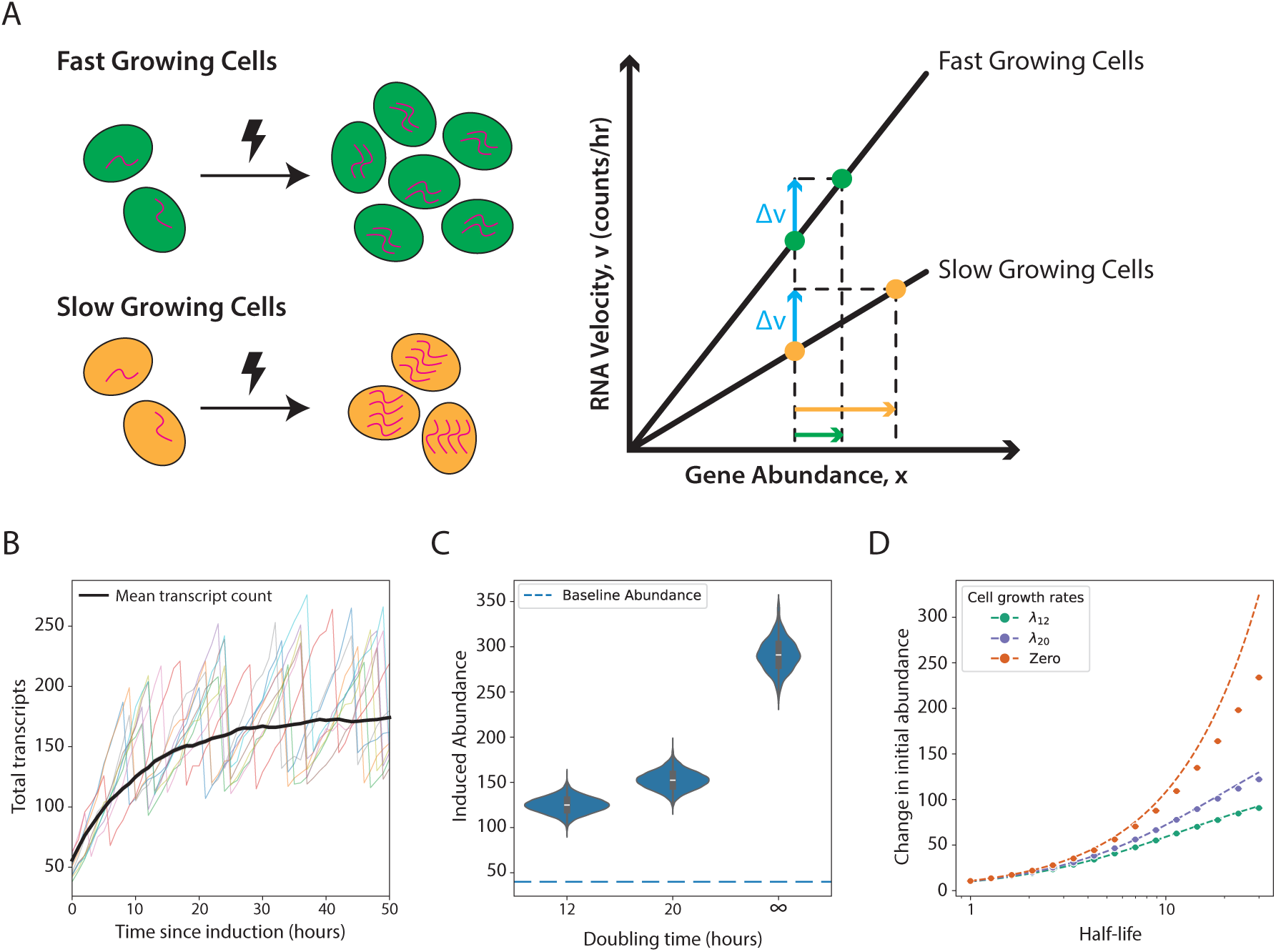
Growth rate impacts induction of genes. (A) Diagram depicting the effect of induction on two populations of cells with different growth rates (left). Illustration of how a constant change in RNA velocity (Δ*v*, blue) results in different changes in gene abundance for fast (green) and slow (orange) growing cells (right). Note that the diagram illustrates the intuition of this effect, but the real change in gene abundance is slightly less than that depicted on the graph (see Methods). (B) Simulated time course of transcript abundance for a representative gene following induction in a cell with a 12-hour doubling time. The black line denotes the mean abundance across simulated cells, and the low-opacity traces depict individual cell trajectories. (C) Violin plots depicting volume-normalized transcript abundances for the gene in (B) after cells have achieved steady state. Volume normalization enables direct comparison with cells simulated under no-growth conditions. Theoretical expectations (lines) and simulated values (points) for the volume-normalized induction size across genes of different half-lives. (D) Change in abundance (y-axis) after induction and allowing for cells to reach steady state for genes with varying half-lives (x-axis), and in cells with different growth rates (colored lines).

